# MAOMAO: An Ontology-Guided Fair Resource for Harmonized Peptide Toxicity Data

**DOI:** 10.64898/2026.07.29.741655

**Authors:** Nicole Soto-Garcia, Roberto Uribe-Paredes, Leandro Murgas, Karen Oróstica, Jorge González-Puelma, Marcelo Navarrete, Frederic Cadet, David Medina-Ortiz

## Abstract

Peptide toxicity is a critical safety and developability parameter in peptide discovery and therapeutic development, yet relevant information remains fragmented across databases, literature resources, and curated datasets. Here, we present MAOMAO, an ontology-guided FAIR-oriented resource that integrates and harmonizes peptide toxicity data from 54 sources. MAOMAO contains 71,701 unique peptide sequences across seven toxicity-related endpoints, represented as 501,907 sequence endpoint combinations with endpoint-specific evidence states and explicit encoding of unavailable information. The resource combines standardized terminology, a hierarchical toxicity vocabulary, evidence-aware state resolution, provenance-aware metadata, 41 physicochemical descriptors, 10 protein language model representations, and a one-hot baseline. It provides endpoint-specific benchmark partitions across splitting strategies and random seeds, reusable with numerical representations. MAOMAO establishes a reusable framework for peptide toxicity research and data-driven toxicology.

## 1 Introduction

Therapeutic peptides have emerged as a rapidly expanding class of biomolecules with applications spanning antimicrobial, antiviral, anticancer, immunomodulatory, metabolic, and diagnostic settings (Goles et al., 2024). Their high target specificity, favorable safety profiles, and growing clinical success have positioned them as attractive alternatives or complements to traditional small-molecule therapeutics (Wang et al., 2022). The number of experimentally characterized bioactive peptides has therefore increased substantially during the last decade (Xiao et al., 2025), supporting the development of specialized databases and computational resources for peptide discovery, functional annotation, and rational design (Cabas-Mora et al., 2024; Ma et al., 2025; Singh et al., 2026; Wang et al., 2016). Despite these advances, toxicity remains a major limitation to peptide development and clinical translation, constraining candidate selection during early-stage screening and optimization (Bahar et al., 2025).

Peptide toxicity encompasses a broad spectrum of adverse biological effects that arise from the interaction between peptides and biological systems. Most toxicity phenotypes are routinely evaluated during peptide development including cytotoxicity, hemolysis, neurotoxicity, embryotoxicity, and ichthyotoxicity (Moreno-Vargas and Prada-Gracia, 2024; Rathore et al., 2025a). These toxicity endpoints have direct implications for efficacy, safety, dosage, and regulatory approval. Toxicity annotations have consequently accumulated across public databases (Singh et al., 2026; Minkiewicz et al., 2019; Pirtskhalava et al., 2021), literature-derived resources (Rathore et al., 2024a,b, 2025b; Wang et al., 2026), supplementary datasets (Karasev et al., 2024; Bhatnagar et al., 2024; Perveen et al., 2023), and experimental studies (Timmons and Hewage, 2020; Yang and Xu, 2024). However, these resources remain fragmented, employ heterogeneous terminology and annotation criteria, and provide variable levels of metadata and evidence quality (Ojeda et al., 2026). Their development for distinct biological objectives further complicates integration, reproducibility, and large-scale analysis (Medina-Ortiz et al., 2026).

The growing availability of toxicity data creates opportunities for computational toxicology, data-driven peptide engineering, and large-scale biological analyses (Soto-Garcia et al., 2026). Their utility, however, depends on the quality, consistency, and transparency of the underlying data (Lones, 2024; Gebru et al., 2022). Annotation ambiguity, conflicting evidence, heterogeneous endpoint definitions, incomplete metadata, and limited provenance tracking can reduce reproducibility and complicate downstream reuse (Al Shareef et al., 2025). The FAIR principles—Findability, Accessibility, Interoperability, and Reusability—provide a framework for organizing and distributing scientific data to support discovery, integration, reproducibility, and long-term reuse (Perez-Riverol et al., 2025). Nevertheless, a standardized framework that harmonizes heterogeneous peptide-toxicity annotations while preserving evidence, provenance, and metadata remains largely unavailable (Ojeda et al., 2026). FAIR-oriented resources tailored to both biological interpretation and computational reuse are therefore still needed (Wise et al., 2019; Leipzig et al., 2021).

To address these limitations, we developed MAOMAO (Metadata-Aware Ontology for Multi-source Annotation Organization), an ontology-guided resource integrating peptide-toxicity information from public databases, literature-derived resources, and curated datasets. MAOMAO standardizes toxicity terminology, preserves source-level provenance, distinguishes uncertainty-related evidence states, and organizes annotations within a unified hierarchical framework. The resource also provides harmonized metadata, descriptor-based representations, protein language model embeddings, benchmark resources, and supporting documentation. These components establish a FAIR-oriented and reusable foundation for biological interpretation, computational toxicology, and predictive modeling of peptide toxicity.

## 2 Materials and Methods

The workflow used to construct MAOMAO is summarized in Figure **1**A. Toxicity-related information from heterogeneous sources was preprocessed, harmonized, mapped to a unified toxicity ontology, evaluated for annotation conflicts and evidence quality, enriched with standardized metadata, and consolidated into the final resource. To support dissemination, computational reuse, and long-term maintenance, MAOMAO was organized into a modular architecture comprising complementary resource layers (Figure **1**B).

**Figure 1:**
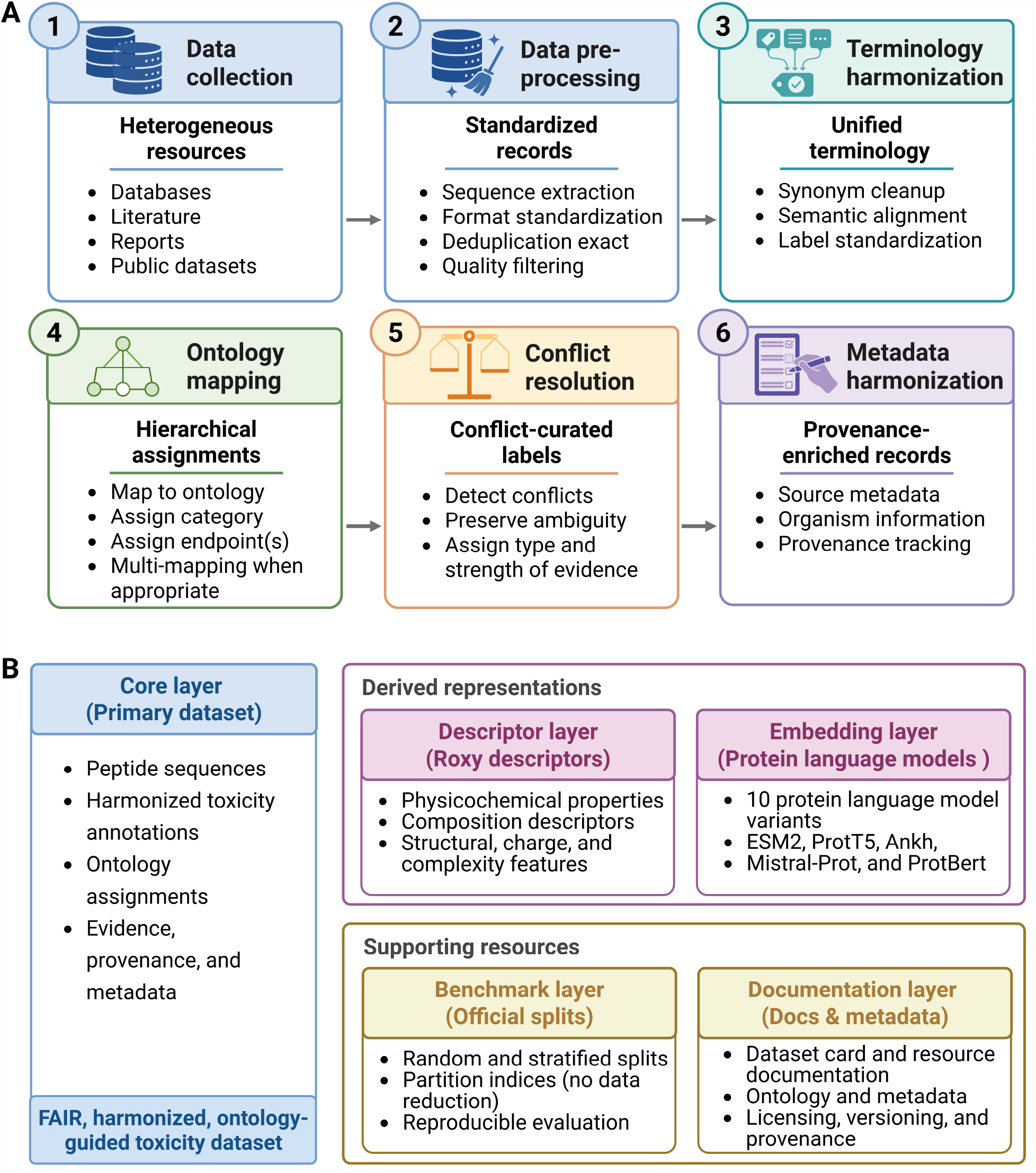
Overview of the MAOMAO construction workflow and resource architecture. (A) Ontology-guided harmonization workflow used to construct the resource. Toxicity-related information collected from public databases, literature-derived datasets, and curated resources undergoes preprocessing, terminology harmonization, ontology mapping, evidence integration, and metadata harmonization to generate a standardized and provenance-aware toxicity resource. (B) Modular architecture of MAOMAO. The resource is organized into complementary layers comprising harmonized toxicity annotations and metadata (Core Layer), sequence-derived descriptors (Descriptor Layer), protein language model embeddings (Embedding Layer), benchmark resources (Benchmark Layer), and supporting documentation (Documentation Layer). Together, these components establish a FAIR-oriented and reusable resource for peptide toxicity research.

### 2.1 Data collection

MAOMAO was constructed by integrating toxicity-related information from publicly available databases, literature-derived resources, supplementary datasets, and manually curated repositories. Candidate resources were identified through a structured exploration of the peptide toxicity landscape using scientific search engines, bibliographic repositories, specialized peptide databases, review articles, and citation tracking (Asim et al., 2025; Mariano et al., 2017). Searches combined general terms, including *peptide toxicity, toxic peptides, toxic peptide databases*, and *toxic peptide predictive models*, with endpoint-specific queries for *hemolytic peptides, cytotoxic peptides, cytolytic peptides, neurotoxic peptides, embryotoxic peptides*, and *ichthyotoxic peptides*. Additional resources were identified from toxicity prediction tools, benchmark datasets, supplementary materials, associated repositories, and specialized peptide resources. Searches were conducted using Google Scholar and PubMed (Fiorini et al., 2017) between June 2025 and June 2026 and covered resources available through June 2026.

Candidate resources were evaluated according to predefined eligibility criteria, including relevance to at least one MAOMAO toxicity endpoint, availability of peptide sequences and corresponding toxicity annotations, sufficient provenance information, and the possibility of programmatic or manual retrieval. Resources were excluded when the underlying dataset could not be retrieved through the reported publication, supplementary material, repository, or web server, or when the available material lacked sequences suitable for sequence-level integration. This included resources containing only numerical representations, peptide names, or functional annotations without the corresponding sequences. Candidate-source screening, eligibility decisions, integrated and excluded resources, and source-specific exclusion reasons are detailed in Supplementary Section S1.

Data acquisition combined direct database downloads, repository and supplementary-file retrieval, and manual extraction when required. Retrieved records were converted into a standardized intermediate representation, preserving peptide sequences, toxicity annotations, source identifiers, literature references, and available metadata. Source-level quality control verified the reported access route, the availability of primary sequences and interpretable toxicity annotations, and the associated publication or repository. Records lacking peptide sequences, containing incomplete toxicity annotations, or presenting irrecoverable formatting inconsistencies were excluded. Retained records were then forwarded to preprocessing, terminology harmonization, ontology mapping, metadata harmonization, and evidence integration.

### 2.2 Data preprocessing, terminology harmonization, and ontology mapping

Following data collection, records underwent standardized preprocessing to support integration across heterogeneous resources (Figure **1**A). This stage included sequence validation, format standardization, annotation parsing, identifier normalization, and quality control. The final resource was restricted to peptides containing the 20 canonical amino acids and lengths between 5 and 70 residues (Xiao et al., 2025; Quiroz et al., 2021). Recoverable formatting inconsistencies were corrected, and exact duplicates within each source were removed. Identical sequences reported by different sources were retained to preserve provenance and support evidence integration. Detailed filtering and sequence-processing criteria are provided in Supplementary Section S1.

To prevent terminology inconsistencies during cross-source integration, toxicity-related annotations were first examined for alternative spellings, synonyms, endpoint-specific descriptions, and source-dependent nomenclature referring to equivalent biological concepts. A terminology harmonization framework was then applied to map these variants to a standardized vocabulary while preserving their original biological meaning. For example, *haemolytic* was normalized to *hemolytic*, and *cytolytic* to *cytolysis*. Endpoint definitions and harmonization rules are provided in Supplementary Section S2.

Harmonized annotations were mapped to a hierarchical controlled vocabulary, hereafter referred to as the MAOMAO toxicity ontology, centered on the root category Toxic. The ontology comprises four primary classes, namely Cytotoxic, Neurotoxic, Embryotoxic, and Ichthyotoxic. The Cytotoxic branch is further divided into the Hemolytic and Cytolysis subclasses. Hemolytic denotes toxicity associated with erythrocyte membrane disruption and lysis, whereas Cytolysis captures cell lysis or membrane-disruptive activity without a specified cell type or biological context (Riss et al., 2016; Nygaard et al., 2025). The vocabulary was restricted to distinct toxicity concepts supported by retrievable sequence-level evidence. Annotations were assigned to one or more categories using predefined rules, with complete definitions, mappings, and assignment criteria provided in Supplementary Section S2.

### 2.3 Conflict resolution and evidence management

Following data collection, harmonization, and ontology mapping, annotations associated with identical peptide sequences were aggregated across sources (Figure **1**A). Because the same sequence could be reported with different toxicity annotations, an evidence-management framework was applied to classify each peptide– endpoint pair as positive, negative, ambiguous, unlabeled, or no information. Positive states represented direct or hierarchy-supported toxicity evidence, whereas negative states corresponded to direct endpoint-specific non-toxic evidence. Ambiguous states captured conflicting or unresolved annotations. Unlabeled states denoted sequences retained in endpoint-specific sources without an explicit class assignment, while no information indicated the absence of evidence for that endpoint.

Negative evidence was further classified as strong or weak according to the supporting information available in the original source. Strong negatives were explicitly supported by experimental evidence, whereas weak negatives originated from randomly selected negative sets or lacked confirmation of the absence of toxicity annotations. Evidence strength was retained in the metadata and provenance records, enabling downstream analyses to account for confidence, uncertainty, and conflicting evidence. Definitions, assignment criteria, and conflict-resolution rules are provided in Supplementary Section S2.

### 2.4 Metadata harmonization and provenance tracking

To support transparency, interoperability, and long-term reuse, source- and endpoint-level metadata were normalized into a unified schema. Harmonized fields included source identifiers, literature references, organism information, annotation provenance, retrieval dates, file formats, and version information when available. Missing values were explicitly preserved without inference. The schema also retained original source labels, ontology mappings, evidence categories, publication references, and resource-specific annotations, providing a standardized representation of the integrated biological and technical information.

Provenance was maintained by linking harmonized annotations to their source-specific processed datasets, endpoint-integration outputs, and metadata records. The sequence-level pivot provides the harmonized access layer, while provenance records, the ambiguity-support table, and the hierarchy-audit table document evidence aggregation, conflict resolution, and hierarchy-derived changes. These files allow each sequence– endpoint state to be traced to its supporting sources and processing decisions. Metadata fields, controlled vocabularies, and data types are detailed in Supplementary Section S3.

### 2.5 Resource architecture

To support reproducibility, computational reuse, and long-term maintenance, MAOMAO was organized into complementary functional layers (Figure **1**B). This modular architecture separates harmonized biological data, computational representations, benchmark resources, and supporting documentation while preserving interoperability across the resource.

The Core Layer contains harmonized peptide sequences, toxicity annotations, ontology mappings, evidence classifications, provenance information, and metadata. Descriptor-based representations were generated using Roxy (KrenAI-Lab, 2026b), whereas embedding-based representations were obtained with Sylphy (KrenAI-Lab, 2026c) using publicly available protein language models.

Benchmark resources were generated using partitioning strategies implemented through BioSieve (KrenAI-Lab, 2026a). The Documentation Layer contains ontology specifications, metadata schemas, resource descriptions, licensing information, and versioning records. Supported representations, benchmark resources, resource organization, and file structures are detailed in Supplementary Sections S4–S5.

### 2.6 FAIR implementation

FAIR principles guided the design and construction of MAOMAO to support transparency, reproducibility, interoperability, and long-term sustainability. FAIR-oriented practices were incorporated across data collection, terminology harmonization, ontology development, evidence management, metadata integration, and resource organization.

Interoperability and reusability were supported through standardized toxicity terminology, ontology-guided endpoint mapping, harmonized metadata schemas, explicit evidence categorization, provenance tracking, and version-aware organization. These components enable heterogeneous toxicity annotations to be integrated while preserving the original information associated with each record.

Findability, accessibility, and computational reuse were supported by distributing resource components in machine-readable formats and using standardized identifiers linking sequences, annotations, metadata, ontology terms, and evidence categories. Harmonization procedures, ontology definitions, evidence-management rules, metadata specifications, and generation workflows were also documented to support validation, updates, and downstream use. Resource availability, file organization, versioning, and supporting documentation are detailed in Supplementary Section S5.

## 3 Resource Description

### 3.1 Dataset overview

An overview of MAOMAO is presented in Figure **2**A. The resource integrates toxicity-related information from 54 publicly available databases, literature-derived resources, supplementary datasets, and manually curated repositories. Of 1,289,119 non-deduplicated source-level records initially collected, 850,642 were retained after source-specific preprocessing, representing a reduction of approximately 34.0%. Removed records included duplicates, incomplete entries, formatting inconsistencies, and annotations that did not satisfy the inclusion criteria.

**Figure 2:**
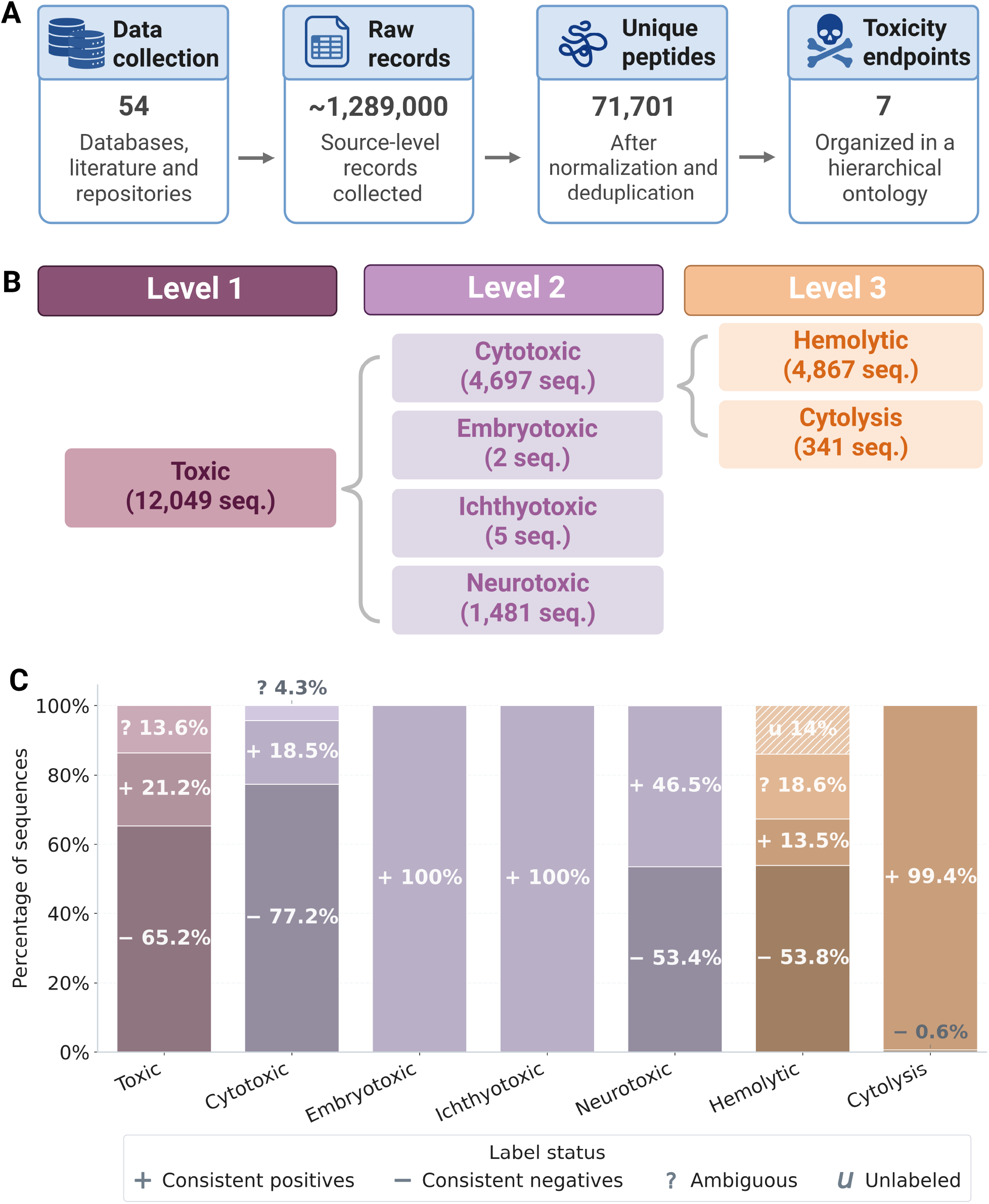
Organization, composition, and coverage of the MAOMAO resource. (A) General overview of the resource, including the number of integrated sources, source-level records, unique peptide sequences, and toxicity-related endpoints. (B) Hierarchical organization of the toxicity endpoints included in MAOMAO. Numbers indicate the number of unique peptide sequences with positive evidence for each end-point after hierarchy application. (C) Composition of sequence–endpoint states across evidence categories, including positive, negative, ambiguous, and unlabeled states, illustrating the distribution of available evidence for each toxicity endpoint.

Following terminology harmonization, ontology mapping, cross-source evidence aggregation, state resolution, and quality control, the final release comprised 71,701 unique peptide sequences across seven toxicity-related endpoints. These were organized into 501,907 sequence–endpoint combinations, together with standardized metadata and provenance records linking annotations, ontology assignments, evidence classifications, and source-specific information.

Publication references, repository or server information, download dates, file formats, source types, and resource behavior were documented for all 54 sources. Annotation content and last-update dates were available for 53 sources (98.15%), whereas explicit licensing information was available for 13 sources (24.07%). Missing metadata were preserved without inference. These components provide a harmonized resource for organizing, disseminating, and computationally analyzing peptide toxicity information.

### 3.2 Toxicity ontology and endpoint organization

The ontology-guided organization of toxicity annotations is presented in Figure **2**B. The root category Toxic contains 56,947 peptide–endpoint states, including 12,049 positives, and comprises four primary classes, namely Cytotoxic (25,453 states; 4,697 positive), Neurotoxic (3,183; 1,481 positive), Embryotoxic (2 positive), and Ichthyotoxic (5 positive). The Cytotoxic branch is further divided into Hemolytic (35,958 states; 4,867 positive) and Cytolysis (343; 341 positive), preserving more specific biological contexts. These categories define the endpoint structure used throughout MAOMAO.

Evidence-state distributions are summarized in Figure **2**C. The final annotation table contains 23,442 positive, 77,843 negative, 15,562 ambiguous, and 5,044 unlabeled peptide–endpoint states. Combinations lacking endpoint-specific evidence were encoded separately as no information and were not considered unlabeled. Toxic contained the largest number of positive states (12,049), whereas *Hemolytic* had the highest proportion of ambiguous states (18.6%) and was the only endpoint containing unlabeled states, representing 14.0% of its available annotations.

Across the six non-root endpoints, 4,510 of the 6,774 peptides with at least one positive annotation were assigned to multiple toxicity categories. This corresponds to 66.58% of positive peptides across these end-points and 6.3% of all MAOMAO sequences. Percentages were calculated relative to the positive sequences assigned to each row endpoint and should therefore be interpreted directionally. The largest overlap occurred between *Cytotoxic* and *Hemolytic*, with 4,279 shared sequences, representing 91.1% of cytotoxic-positive and 87.9% of hemolytic-positive sequences. Complete pairwise counts and directional percentages are provided in Supplementary Section S2.

### 3.3 Resource contents

The MAOMAO release follows the modular architecture presented in Figure **1**B and comprises complementary layers supporting data exploration, computational analysis, benchmarking, and long-term reuse. The release contains more than 8,500 files distributed across harmonized annotations, numerical representations, benchmark resources, metadata, ontology definitions, and documentation.

The Core Layer contains 501,907 harmonized peptide–endpoint combinations corresponding to 71,701 unique sequences, together with ontology assignments, evidence classifications, provenance, and standardized meta-data. The Descriptor Layer provides 41 numerical features covering sequence length, amino acid and residue-class composition, hydrophobicity, structural propensity, charge-related properties, hydrogen-bonding capacity, and sequence complexity. The Embedding Layer includes 10 model-specific matrices generated from protein language models and a one-hot baseline for downstream computational analyses.

The Benchmark Layer stores 240 endpoint–seed–strategy configurations from random and stratified five-fold partitioning of four eligible endpoints—Toxic, Cytotoxic, Hemolytic, and Neurotoxic—using 30 random seeds. These yield 1,200 stored folds containing sequence identifiers and binary labels, which can be joined with 11 Embedding Layer representations to reconstruct 2,640 representation-specific benchmark configurations and 13,200 fold instances. Parameters, reports, summaries, completion records, and execution logs are also included.

The Documentation Layer includes ontology definitions, metadata schemas, resource descriptions, version information, and licensing documentation. Files are distributed in machine-readable formats, including CSV, JSON, and YAML. Detailed descriptions of the resource components and inventories are provided in Supplementary Sections S4–S5.

## 4 Data Value and Reuse Potential

MAOMAO was designed as a reusable resource for peptide toxicity research, providing a harmonized collection of toxicity annotations, ontology mappings, evidence classifications, metadata records, numerical representations, and benchmark resources. By integrating heterogeneous toxicity information into a unified framework, the resource reduces challenges in data acquisition, terminology inconsistencies, annotation ambiguity, and dataset construction, enabling researchers to focus on downstream biological and computational analyses.

MAOMAO supports predictive toxicology. The resource provides standardized toxicity endpoints, evidence-aware annotations, and benchmark partitions that can be used to develop, compare, and validate machine learning models for toxicity prediction. The availability of descriptor-based representations, protein language model embeddings, and predefined dataset partitions facilitates reproducible benchmarking and comparative evaluation of computational approaches.

Beyond supervised learning, MAOMAO supports emerging machine learning paradigms, including positive– unlabeled learning, weak supervision, uncertainty-aware modeling, representation learning, and explainable artificial intelligence. The explicit preservation of annotation provenance, evidence categories, and ontology mappings enables the development of models that account for annotation uncertainty while supporting mechanistic interpretation, model transparency, and explainability analyses.

The resource enables data-centric investigations of peptide toxicity, including toxicity landscape characterization, endpoint co-occurrence analyses, toxicity-aware peptide discovery, and safety-oriented peptide design. The harmonized annotation structure facilitates comparative analyses across toxicity endpoints and supports the development of multi-task and multi-endpoint predictive frameworks that leverage relationships between toxicity categories.

Despite this reuse potential, endpoint coverage depends on the availability of retrievable sequence-level evidence across the curated sources, resulting in class imbalance and limited representation of some endpoints, particularly Cytolysis, Embryotoxic, and Ichthyotoxic. Metadata completeness and evidence strength also vary across sources; although provenance, conflicting annotations, and evidence categories were preserved, this heterogeneity may affect downstream analyses. Finally, the predefined benchmark partitions rely on random and stratified splitting without homology- or similarity-based control and should therefore not be interpreted as leakage-controlled evaluation datasets.

Future releases will address these limitations through the incorporation of newly available toxicity datasets, expanded endpoint coverage, metadata enrichment, updated numerical representations, and additional benchmark resources. The modular architecture of MAOMAO enables these extensions while preserving compatibility and traceability across resource versions.

## Code and Data Availability Statement

The complete, versioned MAOMAO resource is archived on Zenodo: https://zenodo.org/communities/kren-ai-lab (DOI: https://doi.org/10.5281/zenodo.21584414).

MAOMAO-original components—including the harmonization framework, controlled vocabulary, metadata schema, evidence-resolution rules, documentation, and derived records—are distributed under the Creative Commons Attribution 4.0 International (CC BY 4.0) license. Third-party sequences, annotations, and content remain governed by their original providers’ terms. The designation *No information* does not grant unrestricted reuse rights.

Code, workflows, notebooks, and environment specifications are available at https://github.com/kren-ai-lab/maomao under the MIT License.

## Author Contributions Statement

NS-G contributed to conceptualization, data collection and curation, resource construction, ontology development, metadata harmonization, validation, visualization, and manuscript preparation; KO, JG-P, MN, and FC to validation, scientific review, and manuscript revision; DM-O and FC to conceptualization, methodology development, project supervision, resource design, validation, and manuscript preparation; and RU-P and LM to manuscript review and editing. All authors reviewed, revised, and approved the final manuscript.

## Acknowledgments

NS-G and DM-O acknowledge funding from FONDECYT Iniciación 11250295. NS-G also acknowledges funding from “Fondos Concursables Postgrado: Fortalecimiento de Tesis y Actividad Formativa Equivalente” (MAG23992). DM-O and RU-P acknowledge support from the Center for Biotechnology and Bioengineering (CeBiB; PIA projects FB0001 and AFB240001, ANID, Chile). PEACCEL is supported by a research program partially cofunded by the European Union and Région Réunion (FEDER 2021–2027). MN’s work is funded by ANID through FONDECYT 1230298.

## AI Statement

As non-native English speakers, the authors used ChatGPT (OpenAI, GPT-5.6 Thinking; accessed July 2026) for English editing, clarity, and consistency. The authors critically reviewed and edited all outputs, verified the final text, and take full responsibility for the manuscript.

