## Supplementary Information for "MAOMAO: An Ontology-Guided Fair Resource for Harmonized Peptide Toxicity Data"

#### Supplementary contents

**S1 Dataset construction and source curation** **2**

#### S1 Dataset construction and source curation

This section outlines how the MAOMAO resource was constructed and refined. It covers the identification, selection, extraction, and harmonization of source datasets, and the procedures applied to resolve duplicate sequences and conflicting annotations prior to further processing.

##### S1.1 Bibliographic search and source identification

Candidate sources were identified through a structured search focused on publicly available resources containing sequence-level information associated with peptide toxicity. The search was conducted between June 2025 and June 2026 using Google Scholar and PubMed, and covered resources available through June 2026. The objective was to identify databases, published datasets, supplementary materials, repositories, and datasets associated with peptide toxicity prediction studies that could contribute sequences and annotations to at least one of the toxicity endpoints represented in MAOMAO. Search terms combined general concepts related to peptide toxicity and toxicity-oriented data resources. Searches were not restricted by publication year, and resources available up to June 2026 were considered. The bibliographic search was complemented by inspection of review articles, specialized peptide databases, publisher pages, article supplementary materials, associated repositories, and resources linked to peptide toxicity prediction tools and benchmark datasets. Citation tracking was also performed using references and dataset descriptions from relevant studies to identify earlier datasets, database versions, and primary sources that had been reused, reformatted, or expanded in subsequent publications. Table **S1** summarizes the main search dimensions used during candidate-source identification. Priority was given to resources that provided primary peptide sequences together with toxicity annotations or labels that could be interpreted and mapped to the MAOMAO terminology.

**Supplementary Table S1**: Search strategy used to identify candidate sources for MAOMAO construction.

| Search dimension | Terms or sources | Purpose |
|---|---|---|
| Bibliographic search | Google Scholar and PubMed | Identify publications, databases, prediction studies, and publicly available datasets containing peptide-toxicity information. |
| General toxicity concepts | peptide toxicity, toxic peptides, toxic peptide databases, and toxic peptide predictive models | Identify general toxicity resources and datasets used for peptide-toxicity research or predictive modeling. |
| Endpoint-specific search | hemolytic peptides, cytotoxic peptides, cytolytic peptides, neurotoxic peptides, embryotoxic peptides, and ichthyotoxic peptides | Identify resources focused on the specific toxicity endpoints represented in MAOMAO. |

*Continued on next page*

**Supplementary Table S1** – *Continued from previous page*

| Search dimension | Terms or sources | Purpose |
|---|---|---|
| Prediction and benchmark resources | Peptide-toxicity prediction tools, model publications, and benchmark datasets | Identify sequence datasets used to train, validate, or compare peptide-toxicity prediction models. |
| Specialized resource inspection | Peptide databases, review articles, publisher pages, and manually curated repositories | Identify candidate resources not recovered directly through the bibliographic search. |
| Data availability screening | Article supplementary materials, database downloads, associated repositories, web servers, and linked code or data repositories | Determine whether primary peptide sequences and their corresponding toxicity annotations could be retrieved. |
| Citation tracking | Reference lists, dataset descriptions, and cited sources from relevant peptide-toxicity studies | Identify earlier datasets, original data sources, database versions, and resources reused or expanded in subsequent publications. |

Candidate resources were initially recorded independently for each toxicity endpoint under which they were identified. A source associated with more than one endpoint could therefore appear in multiple endpoint-specific searches. For the calculation of global source-selection counts, source names were standardized and repeated resources were counted only once. Distinct database versions were retained as separate sources when they represented independently released or curated datasets.

When a publication, database, or repository provided multiple files, supplementary tables, dataset partitions, or endpoint-specific subsets, these components were treated as part of the same source rather than as independent sources. All retrieved files were retained during the initial compilation step to preserve their provenance and to support subsequent source-level and record-level consistency assessment. Source name, publication or repository, access route, retrieval date, file format, toxicity endpoint, sequence counts, and available annotation information were recorded before preprocessing, harmonization, and cross-source integration.

#### S1.2 Inclusion and exclusion criteria

Candidate sources were considered eligible for sequence-level curation when they met all of the following criteria:

(i) the source contained information relevant to at least one toxicity endpoint represented in MAOMAO;

(ii) peptide sequences were available from the database, publication, supplementary material, web server, or an associated repository;

(iii) the source provided toxicity annotations, labels, activity descriptions, or dataset-class definitions that could be interpreted and mapped to the MAOMAO terminology;

(iv) the underlying data could be retrieved through a documented access route using programmatic or manual procedures.

(v) sufficient provenance information was available to associate the retrieved records with the corresponding database, publication, dataset, repository, or other documented source.

Candidate sources were excluded when the underlying dataset could not be retrieved through the reported publication, supplementary material, database, repository, or web server. Sources were also excluded when the accessible material did not provide the peptide sequences required for sequence-level integration. This category included resources containing only numerical representations, peptide names, Gene Ontology annotations, or other functional metadata without the corresponding amino-acid sequences.

Source-level exclusion was distinguished from record-level filtering. A source was considered included when at least part of its sequence-level dataset met the eligibility criteria and could be retrieved. After source inclusion, individual records could still be removed during preprocessing when they lacked peptide sequences, contained incomplete or uninterpretable toxicity annotations, presented irrecoverable formatting inconsistencies, or did not satisfy the sequence-quality criteria applied during harmonization.

#### S1.3 Source screening and selection outcome

After standardization of source names, 76 unique candidate sources were identified and assessed for eligibility. Of these, 54 met the inclusion criteria and were retained for sequence-level curation and integration, whereas 22 were excluded. Among the excluded sources, 18 could not be incorporated because the corresponding datasets were either unavailable through the reported access routes or no data-access route was provided. The remaining 4 did not provide usable peptide sequences: 2 contained only numerical representations, 1 listed peptide names without the corresponding sequences, and 1 reported peptide names and Gene Ontology annotations without the corresponding sequences.

Table **S2** summarizes the source-identification and eligibility-assessment process, including the number of unique candidate sources evaluated, the principal reasons for exclusion, and the number of sources retained for MAOMAO construction.

**Supplementary Table S2**: Summary of candidate-source identification, eligibility assessment, and selection for MAOMAO construction. Counts correspond to unique sources after source-name standardization.

| Selection stage | Sources | Description |
|---|---|---|
| Candidate sources identified | 76 | Unique candidate resources identified through bibliographic searches, resource inspection, and citation tracking. |

*Continued on next page*

**Supplementary Table S2** – *Continued from previous page*

| Selection stage | Sources | Description |
|---|---|---|
| Candidate sources evaluated | 76 | Resources assessed according to toxicity relevance, sequence availability, annotation accessibility, data retrievability, and provenance. |
| Sources excluded | 22 | Resources that did not satisfy the source-level eligibility criteria. |
| Underlying data unavailable | 18 | The corresponding dataset could not be retrieved through the reported publication, supplementary material, database, repository, or web server, or no data-access route was provided. |
| No usable primary sequences | 4 | The accessible resources provided no sequences, only numerical representations, or names and functional annotations without the corresponding amino-acid sequences. |
| Sources integrated | 54 | Eligible resources retained for sequence extraction, preprocessing, harmonization, ontology mapping, and evidence integration. |

These counts represent unique sources rather than source–endpoint associations, because a single resource could contribute information to more than one toxicity endpoint. The excluded candidate sources and their source-specific reasons for exclusion are reported in Table **S3**, whereas the endpoint coverage, retained sequence counts, and annotation categories of the 54 integrated sources are summarized in Table **S4**.

#### S1.4 Excluded candidate sources

Not all candidate resources identified during source discovery could be incorporated into MAOMAO. Table **S3** lists the 22 sources that did not meet the source-level eligibility criteria and summarizes the corresponding reasons for exclusion.

**Supplementary Table S3**: Candidate peptide-toxicity sources excluded during source-level eligibility assessment, grouped according to the primary reason for exclusion.

| Reason for exclusion | Sources | Candidate sources |
|---|---|---|
| Dataset unavailable or no usable data-access route | 18 | Toropov et al. [Toropov et al., 2025]; AMP-D3 (HemolyticPredictor) [Ouyang et al., 2025]; Kumar et al. [Kumar et al., 2025]; AmpLyze [Qiu et al., 2025]; Castillo-Mendieta et al. [Castillo-Mendieta et al., 2024b]; Ansari et al. [Ansari and White, 2023a]; MoFormer [Wang et al., 2024a]; MQSS model [Castillo-Mendieta et al., 2024a]; Ansari et al. (2023) [Ansari and White, 2023b]; HemoPred [Win et al., 2017]; Wan et al. [Wan et al., 2023]; PEP-PREDNa+ [Herrera-Bravo et al., 2022]; Neu_LR [Zhu et al., 2021]; M3-CAD [Wang et al., 2024b]; PepNet [Han et al., 2024]; ToxMVA [Shi et al., 2022]; ATSE [Wei et al., 2021]; and Khabbaz et al. [Khabbaz et al., 2021] |
| Only numerical representations available | 2 | AIPAMPDS (HemoRisk-Estimator) [Li et al., 2025] and ToxDL [Pan et al., 2020]. |
| Only peptide names and/or functional annotations available | 2 | PlantPepDB [Das et al., 2020] and NNTox [Jain and Kihara, 2019]. |

For PlantPepDB, the accessible material provided only peptide names and did not include the corresponding sequences required for sequence-level integration. NNTox provided peptide names and Gene Ontology annotations, but not the corresponding amino-acid sequences.

These exclusions were applied at the source level before record extraction and were therefore distinct from the subsequent record-level filtering of entries with missing sequences, uninterpretable toxicity annotations, or irrecoverable formatting inconsistencies within the 54 eligible sources.

#### S1.5 Source datasets used for peptide toxicity data curation

This subsection presents the source datasets that were kept for building the toxic peptides dataset following the bibliographic search and eligibility screening. Table **S4** shows, for each source, the toxicity effects covered, the number of records retained after source-specific processing, and the distribution of positive, negative, unlabeled, and other entries reported or derived from each dataset. These figures reflect source-level data before cross-source consolidation, global terminology harmonization, evidence-state resolution, and hierarchy application.

**Supplementary Table S4**: Overview of source datasets used for peptide toxicity data curation. Retained values are non-deduplicated source-level record counts after source-specific processing. Annotation-category counts describe the retained records before cross-source consolidation, evidence-state resolution, and hierarchy application. The *Other* column denotes source-specific records outside the positive, negative, and unlabeled categories, such as regression records. Across the 54 source metadata records, 1,289,119 raw records were reported and 850,642 records were retained.

| Source | Toxicity endpoint(s) | Retained records | Positive | Negative | Unlabeled | Other |
|---|---|---|---|---|---|---|
| Abdelbaky et al. [Abdelbaky et al., 2024] | Hemolytic; toxic | 5851 | 1983 | 3868 | 0 | 0 |
| Almotairi et al. [Almotairi et al., 2024] | Hemolytic; toxic | 5400 | 1907 | 3493 | 0 | 0 |
| AMPDB v1 [Mondal et al., 2023] | Toxic; cytolytic; cytotoxic; hemolytic; ichthyotoxic; other toxicity-related effects | 5230 | 5230 | 0 | 0 | 0 |
| AMPDeep [Salem et al., 2022] | Hemolytic; toxic | 16191 | 9449 | 6742 | 0 | 0 |
| Bhatnagar et al. [Bhatnagar et al., 2024] | Hemolytic; toxic | 756 | 167 | 589 | 0 | 0 |
| BIOPEP-UWM [Minkiewicz et al., 2019] | Hemolytic; toxic; cytotoxic; embryotoxic | 89 | 89 | 0 | 0 | 0 |
| BiToxNet [Wang et al., 2026] | Neurotoxic; toxic | 7780 | 2399 | 5381 | 0 | 0 |
| CAPTP [Jiao et al., 2024] | Toxic | 7513 | 2138 | 5375 | 0 | 0 |
| CICERON [Bizzotto et al., 2024] | Hemolytic; toxic; embryotoxic; cytotoxic | 82 | 82 | 0 | 0 | 0 |
| ConsAMPHemo [Xie et al., 2025] | Hemolytic; toxic | 5841 | 2246 | 3595 | 0 | 0 |
| csm-toxin [Morozov et al., 2023] | Toxic | 221634 | 7392 | 214242 | 0 | 0 |
| DRAMP [Ma et al., 2025] | Toxic; hemolytic; cytolytic; cytotoxic | 3225 | 530 | 2695 | 0 | 0 |
| HAPPENN [Timmons and Hewage, 2020] | Hemolytic; toxic | 3532 | 1456 | 2076 | 0 | 0 |
| HemoDL [Yang and Xu, 2024] | Hemolytic; toxic | 5437 | 2369 | 3068 | 0 | 0 |
| HemoFuse [Zhao et al., 2024] | Hemolytic; toxic | 5437 | 2369 | 3068 | 0 | 0 |
| Hemolytic-Pred [Perveen et al., 2023] | Hemolytic; toxic | 1235 | 0 | 0 | 1235 | 0 |
| Hemolytik [Gautam et al., 2014] | Hemolytic; toxic | 2471 | 2471 | 0 | 0 | 0 |
| hemolytik 2.0 [Singh et al., 2025] | Hemolytic; toxic | 4960 | 4960 | 0 | 0 | 0 |
| hemonet [Yaseen et al., 2021] | Hemolytic; toxic | 3308 | 1422 | 1851 | 35 | 0 |
| HemoPI [Chaudhary et al., 2016] | Hemolytic; toxic | 2191 | 910 | 1281 | 0 | 0 |
| HemoPI2.0 [Rathore et al., 2025b] | Hemolytic; toxic | 1926 | 891 | 1035 | 0 | 0 |
| HEPAD [Chen et al., 2025] | Hemolytic; toxic | 2996 | 910 | 2086 | 0 | 0 |
| HLPpred-Fuse [Hasan et al., 2020] | Hemolytic; toxic | 4365 | 1360 | 3005 | 0 | 0 |
| HMAMP-main [Wang et al., 2025a] | Hemolytic; toxic | 1656 | 552 | 552 | 552 | 0 |
| HyPepTox-Fuse [Tran et al., 2025] | Toxic | 11036 | 5518 | 5518 | 0 | 0 |
| iAMPCN [Xu et al., 2023] | Cytotoxic; hemolytic; toxic | 61592 | 2662 | 58930 | 0 | 0 |
| Karasev et al. [Karasev et al., 2024] | Hemolytic; toxic | 1740 | 0 | 0 | 944 | 796 |
| MultiPep [Grønning et al., 2021] | Hemolytic; toxic; embryotoxic; other toxicity-related effects | 6400 | 6400 | 0 | 0 | 0 |
| Multitox [Sharma et al., 2025] | Neurotoxic; hemolytic; cytotoxic; toxic | 1898 | 1898 | 0 | 0 | 0 |
| NTXpred [Saha and Raghava, 2007] | Neurotoxic; toxic | 916 | 916 | 0 | 0 | 0 |
| NTXpred2 [Rathore et al., 2025a] | Neurotoxic; toxic | 3300 | 1649 | 1651 | 0 | 0 |
| Pep-Lab_db [Terziyski et al., 2023] | Toxic | 249 | 249 | 0 | 0 | 0 |
| peptidereactor [Spänig et al., 2021] | Hemolytic; toxic | 1104 | 522 | 582 | 0 | 0 |
| Peptipedia 2.0 [Cabas-Mora et al., 2024] | Toxic; neurotoxic; hemolytic; cytotoxic | 22455 | 22455 | 0 | 0 | 0 |
| PeptiTox [Wang et al., 2025b] | Toxic | 3864 | 1932 | 1932 | 0 | 0 |
| Plisson et al. [Plisson et al., 2020] | Hemolytic; toxic | 9625 | 1076 | 1275 | 7274 | 0 |
| PLPTP [Gao et al., 2025] | Toxic | 7513 | 2138 | 5375 | 0 | 0 |
| ProToxin [Yang et al., 2025] | Toxic | 252536 | 6954 | 244829 | 753 | 0 |
| QSVM-PEPTIDE [Zhuang et al., 2024] | Hemolytic; toxic | 2195 | 911 | 1284 | 0 | 0 |
| SATPdb [Singh et al., 2016] | Toxic | 4130 | 4130 | 0 | 0 | 0 |
| StrucToxNet [Jiao et al., 2025] | Toxic | 9544 | 2491 | 7053 | 0 | 0 |
| tAMPer [Ebrahimikondori et al., 2024] | Toxic | 5708 | 1790 | 3918 | 0 | 0 |
| ToxDL 2.0 [Zhu et al., 2025] | Toxic | 11434 | 4593 | 6841 | 0 | 0 |
| ToxGIN [Yu et al., 2024] | Toxic | 4410 | 2205 | 2205 | 0 | 0 |
| ToxIBTL [Wei et al., 2022] | Toxic | 14092 | 5698 | 8394 | 0 | 0 |
| ToxinPred [Gupta et al., 2013] | Toxic | 19391 | 2084 | 17307 | 0 | 0 |
| ToxinPred 2.0 [Sharma et al., 2022] | Toxic | 35242 | 8233 | 27009 | 0 | 0 |
| ToxinPred 3.0 [Rathore et al., 2023] | Toxic | 11036 | 5518 | 5518 | 0 | 0 |

*Continued on next page*

**Supplementary Table S4**: Overview of source datasets used for peptide toxicity data curation. Continued.

| Source | Toxicity endpoint(s) | Retained records | Positive | Negative | Unlabeled | Other |
|---|---|---|---|---|---|---|
| ToxiPep [Guan et al., 2025] | Toxic | 9889 | 4938 | 4951 | 0 | 0 |
| ToxMSRC [Zhang et al., 2025] | Toxic | 7513 | 7513 | 0 | 0 | 0 |
| ToxTeller [Wang and Sung, 2024] | Toxic | 4329 | 2078 | 2251 | 0 | 0 |
| TPpred-LE [Lv et al., 2023] | Toxic | 2345 | 2345 | 0 | 0 | 0 |
| UniDL4BioPep [Du et al., 2023] | Toxic | 3864 | 1932 | 1932 | 0 | 0 |
| Zhao et al. [Zhao et al., 2021] | Hemolytic; toxic | 2186 | 1093 | 1093 | 0 | 0 |
| **Total** | — | **850,642** | **165,203** | **673,850** | **10,793** | **796** |

Across all source-level data records, 1,289,119 raw records were reported and 850,642 were retained after source-specific processing, corresponding to an overall reduction of approximately 34.0%. These values represent non-deduplicated source-level totals.

#### S1.6 Source-level parsing and sequence quality control

Following source selection, records obtained from each eligible resource were parsed and converted into a standardized sequence-level representation. Peptide sequences, source-specific toxicity annotations, source identifiers, literature or repository references, and available metadata were preserved whenever possible. Records were excluded when the peptide sequence was missing, when the corresponding toxicity annotation could not be interpreted for subsequent harmonization and mapping to the MAOMAO terminology, or when irrecoverable formatting inconsistencies prevented conversion into a valid sequence–annotation record.

Sequence-level quality control was subsequently applied using predefined residue-composition and length criteria. Peptide sequences were retained when their length ranged from 5 to 70 amino-acid residues, inclusive, and every sequence character belonged to the canonical amino-acid vocabulary: `A, C, D, E, F, G, H, I, K, L, M, N, P, Q, R, S, T, V, W, Y`. Sequences containing at least one residue outside this vocabulary, as well as sequences shorter than 5 or longer than 70 residues, were excluded from downstream integration.

Exact duplicate records occurring within the same source were removed during source-level processing. In contrast, identical peptide sequences reported by different sources were retained together with their source-specific annotations and provenance information. This distinction prevented repeated records within a source from artificially increasing its contribution while preserving evidence reported by different resources.

Conflicting annotations were handled separately during evidence integration. Peptide–endpoint pairs with conflicting or otherwise unresolved evidence across sources were classified as *ambiguous*, while preserving the provenance of the contributing sources.

The quality-control configuration and its outcomes were documented in the corresponding metadata records. Source-level metadata included the numbers of raw, retained, erroneous, and modified sequences. Endpoint-level metadata recorded whether the canonical-residue and length filters were applied, the minimum and maximum accepted sequence lengths, the number of sequences before and after each filter, sequence-length distribution statistics, and the final number of sequences retained for each toxicity endpoint. The record-processing and sequence-quality rules applied during MAOMAO construction are summarized in Table **S5**.

**Supplementary Table S5**: Record-processing, sequence-level quality-control, and annotation-integration rules applied during MAOMAO construction.

| Processing case | Rule applied | Treatment or interpretation |
|---|---|---|
| Missing peptide sequence | No primary amino-acid sequence was available for the record. | The record was excluded from sequence-level integration. |
| Missing or uninterpretable toxicity annotation | The annotation, label, activity description, or class definition could not be interpreted for subsequent terminology harmonization. | The record was excluded from toxicity-annotation integration. |
| Recoverable formatting inconsistency | Formatting differences could be corrected without changing the biological meaning of the sequence or annotation. | The record was standardized and retained. |
| Irrecoverable formatting inconsistency | The record could not be converted into a valid sequence–annotation entry without unsupported interpretation. | The record was excluded during source-level processing. |
| Non-canonical amino-acid residue | At least one character was not included in the canonical vocabulary `A, C, D, E, F, G, H, I, K, L, M, N, P, Q, R, S, T, V, W, Y`. | The sequence was excluded during canonical-residue filtering. |
| Sequence outside the accepted length range | The sequence contained fewer than 5 or more than 70 amino-acid residues. | The sequence was excluded during length filtering. |
| Sequence passing both filters | The sequence contained only canonical amino-acid residues and had a length between 5 and 70 residues, inclusive. | The sequence was retained for downstream harmonization and integration. |
| Exact duplicate within the same source | The same sequence and source-specific annotation record occurred more than once within a source. | Duplicate records were removed during source-level processing. |

*Continued on next page*

**Supplementary Table S5**: Record-processing, sequence-level quality-control, and annotation-integration rules applied during MAOMAO construction. Continued.

| Processing case | Rule applied | Treatment or interpretation |
|---|---|---|
| Identical sequence across different sources | The same peptide sequence was reported by more than one source. | The sequence was retained together with all source-specific annotations and provenance records. |
| Conflicting annotations across sources | The same peptide–endpoint pair was supported by both positive and negative evidence across different sources. | The integrated annotation was classified as *ambiguous*. |

### S2 Terminology harmonization and toxicity ontology

#### S2.1 Endpoint definitions used for ontology-guided harmonization

The MAOMAO toxicity ontology was developed as an operational framework for terminology harmonization and annotation integration rather than as an exhaustive classification of toxicological mechanisms. This approach is consistent with the use of controlled vocabularies and ontology-based annotations to structure and integrate heterogeneous toxicological knowledge [Ives et al., 2017, Mortensen et al., 2022]. The parent–child relationships were defined according to the biological scope of each reported toxicity effect and are summarized in Table **S6**.

The root category Toxic represents generic adverse effects, whereas Cytotoxic, Neurotoxic, Embryotoxic, and Ichthyotoxic represent more specific biological contexts. Hemolytic and Cytolysis were operationally assigned as subclasses of *Cytotoxic* because both describe specific forms of cell damage or membrane disruption, with hemolysis referring specifically to erythrocyte lysis [Spaller et al., 2013, Wimley, 2010]. Endpoints with limited current evidence were retained when they represented distinct toxicity concepts explicitly reported by the curated sources. Their inclusion in the ontology preserves biological and terminological coverage but does not necessarily imply sufficient class representation for supervised binary benchmarking.

**Supplementary Table S6**: Ontology endpoint definitions used for terminology harmonization and endpoint assignment. The table summarizes the standardized toxicity endpoints used in the resource, their position in the toxicity ontology, and the operational interpretation used during annotation harmonization.

| Standard endpoint | Parent class | Operational definition | Representative source terms |
|---|---|---|---|
| *Toxic* | Root | Generic toxicity category used for peptides reported to produce an adverse biological effect, or for sequences with positive evidence in any child toxicity endpoint. This class provides the root level for ontology-guided integration [Sharma et al., 2022]. | toxic; toxicity; toxic peptide; toxin |
| *Cytotoxic* | *Toxic* | Toxicity toward cells or cell-based systems, including viability reduction, membrane damage, or cell death in eukaryotic or mammalian cellular contexts. Positive evidence from cytotoxicity-associated subclasses may be propagated to this parent category [Aslantürk, 2017]. | cytotoxic; cytotoxicity |
| *Hemolytic* | *Cytotoxic* | Toxicity associated with erythrocyte damage or red blood cell lysis. This endpoint is represented as a cytotoxicity-associated subclass because hemolysis reflects cell membrane disruption in erythrocytes [Spaller et al., 2013]. | hemolytic; haemolytic; hemolysis; haemolysis; hemotoxicity |
| *Cytolysis* | *Cytotoxic* | Cell lysis or membrane-disruptive activity reported as a toxicity-related effect. This category was used when the annotation specifically referred to cell lysis rather than general cytotoxicity [Wimley, 2010]. | cytolytic; cytolysis; cell lysis; lytic activity; membranolytic activity |
| *Neurotoxic* | *Toxic* | Toxicity affecting neurons, neural cells, or the nervous system. This class was assigned only when the source annotation explicitly indicated neurotoxicity or toxin-like effects on neural systems [McQueen, 2017]. | neurotoxic; neurotoxicity; neurotoxin |
| *Embryotoxic* | *Toxic* | Toxicity affecting embryos or developmental stages. This endpoint was used for annotations explicitly linked to embryo viability, developmental toxicity, or embryotoxic effects [Baker et al., 2018]. | embryotoxic; embryotoxicity |
| *Ichthyotoxic* | *Toxic* | Toxicity affecting fish or fish-derived biological systems. This endpoint was used when the source annotation explicitly indicated fish toxicity, piscicidal activity, or toxicity in fish models [Bambino and Chu, 2017]. | ichthyotoxic; ichthyotoxicity |

#### S2.2 Evidence-state definitions and conflict resolution

After terminology harmonization, source-specific annotations associated with the same peptide–endpoint pair were aggregated into one mutually exclusive state in the final sequence-level pivot. The evidence model distinguishes direct endpoint evidence, unresolved or conflicting evidence, source-associated records without an explicit class assignment, and the absence of information for a specific endpoint. Table **S7** summarizes the five states used in MAOMAO.

**Supplementary Table S7**: Evidence states used in the final MAOMAO sequence-level pivot.

| Code | State | Assignment and interpretation |
|---|---|---|
| 1 | *Positive* | Direct positive evidence for the endpoint, or positive evidence propagated from a child endpoint when the parent endpoint was not ambiguous. |
| 0 | *Negative* | Direct endpoint-specific evidence supporting the absence of the corresponding toxicity effect. Negative evidence was not propagated through the hierarchy. |
| 2 | *Ambiguous* | Conflicting or otherwise unresolved evidence for the same sequence–endpoint pair. Endpoint-specific ambiguity was retained and could not be overwritten by hierarchical positivity. |
| 3 | *Unlabeled* | The sequence was retained from a source associated with the endpoint, but no explicit positive or negative assignment was available for that record. |
| 999 | *No information* | No annotation was available for that endpoint. This code is distinct from both negative and unlabeled evidence. |

When only positive evidence was available, the integrated state was positive; when only direct negative evidence was available, the state was negative. Peptide–endpoint pairs supported by both positive and negative annotations, or by evidence that could not be resolved without unsupported interpretation, were classified as ambiguous. Unlabeled states were assigned only when a sequence was explicitly retained by an endpoint-associated source without a class assignment. All remaining sequence–endpoint combinations in the complete cross-product were encoded as no information.

Negative evidence was additionally characterized at the provenance level as *strong* or *weak*. Strong negatives corresponded to records for which the absence of toxicity was explicitly supported by experimental or curated evidence. Weak negatives originated from resources in which negative sequences were randomly selected or lacked evidence confirming the absence of the effect. Evidence strength was retained in the supporting metadata and provenance records and was not treated as an additional pivot state.

#### S2.3 Hierarchy propagation rules and audit statistics

Hierarchy application followed a positive-only policy. Positive support could propagate from a more specific endpoint to its parent, whereas negative, ambiguous, unlabeled, and no-information states were never propagated. Direct ambiguity at the parent endpoint blocked hierarchy-derived positivity, thereby preserving unresolved endpoint-specific evidence. The rules applied during final annotation resolution are summarized in Table **S8**.

**Supplementary Table S8**: Terminology-harmonization and hierarchy-propagation rules applied during MAOMAO construction.

| Rule | Source endpoint(s) | Target endpoint | Condition and action |
|---|---|---|---|
| R1 | *Cytotoxic*, *Neurotoxic*, *Embryotoxic*, or *Ichthyotoxic* | *Toxic* | Positive child support set the parent to positive unless *Toxic* contained direct ambiguous evidence. |
| R2 | *Hemolytic* or *Cytolysis* | *Cytotoxic* | Positive child support set the parent to positive unless *Cytotoxic* contained direct ambiguous evidence. |
| R3 | Source terms *cytolysis* and *cytolytic* | *Cytolysis* | Equivalent source terminology was harmonized before evidence aggregation and final state resolution. |

The hierarchy audit records every sequence–endpoint pair with positive child support and indicates whether the parent state was inferred, blocked by ambiguity. Table **S9** summarizes the final audit.

**Supplementary Table S9**: Summary of hierarchy application in the final MAOMAO release.

| Target endpoint | Positive child support | Rows inferred | Blocked by ambiguity |
|---|---|---|---|
| *Cytotoxic* | 5,057 | 4,123 | 620 |
| *Toxic* | 6,774 | 3,614 | 1,146 |
| **Total** | **11,831** | **7,737** | **1,766** |

#### S2.4 Final endpoint-state composition

The final pivot contains 71,701 unique peptide sequences and seven endpoint columns, yielding 501,907 sequence–endpoint combinations. Table **S10** reports the mutually exclusive sequence–endpoint state counts after hierarchy application. All values in the table, including the *Total* row, correspond to sequence–endpoint states. The *Available states* column includes positive, negative, ambiguous, and unlabeled states and excludes code 999.

Across all endpoints, 121,891 sequence–endpoint combinations contained an available evidence state. Of the 15,562 ambiguous sequence–endpoint states, 13,027 unique peptide sequences had at least one ambiguous endpoint, because a single sequence may be ambiguous for more than one endpoint. The remaining 380,016 sequence–endpoint combinations were encoded as no information (999).

**Supplementary Table S10**: Final mutually exclusive evidence-state counts for each MAOMAO toxicity endpoint.

| Endpoint | Positive | Negative | Ambigu-ous | Unlabeled | Available states |
|---|---|---|---|---|---|
| *Toxic* | 12,049 | 37,129 | 7,769 | 0 | 56,947 |
| *Cytotoxic* | 4,697 | 19,662 | 1,094 | 0 | 25,453 |
| *Hemolytic* | 4,867 | 19,350 | 6,697 | 5,044 | 35,958 |
| *Cytolysis* | 341 | 2 | 0 | 0 | 343 |
| *Neurotoxic* | 1,481 | 1,700 | 2 | 0 | 3,183 |
| *Embryotoxic* | 2 | 0 | 0 | 0 | 2 |
| *Ichthyotoxic* | 5 | 0 | 0 | 0 | 5 |
| **Total** | **23,442** | **77,843** | **15,562** | **5,044** | **121,891** |

#### S2.5 Positive endpoint co-annotation analysis

Positive endpoint co-annotation was evaluated across the six non-root toxicity endpoints using the final sequence-level pivot after hierarchy application. The root endpoint *Toxic* was excluded because its positive states include both direct and hierarchy-derived evidence. Only states encoded as positive were considered; negative, ambiguous, unlabeled, and no-information states were excluded.

A total of 6,774 unique peptide sequences had at least one positive annotation across the six evaluated endpoints. Of these, 4,510 were positive for more than one endpoint, corresponding to 66.58% of sequences with at least one positive non-root annotation and 6.3% of all 71,701 unique sequences in MAOMAO.

For each endpoint pair, the number of shared positive sequences was calculated (Table **S11**). Directional percentages were obtained by dividing each pairwise count by the total number of positive sequences assigned to the corresponding row endpoint (Table **S12**). Consequently, reciprocal percentages may differ between endpoint pairs.

**Supplementary Table S11**: Pairwise counts of shared positive peptide sequences across the six non-root MAOMAO toxicity endpoints. Diagonal values correspond to the total number of positive sequences assigned to each endpoint.

| Row endpoint | Cytotoxic | Embryotoxic | Ichthyotoxic | Neurotoxic | Hemolytic | Cytolysis |
|---|---|---|---|---|---|---|
| Cytotoxic | 4,697 | 0 | 3 | 20 | 4,279 | 247 |
| Embryotoxic | 0 | 2 | 0 | 0 | 0 | 0 |
| Ichthyotoxic | 3 | 0 | 5 | 0 | 2 | 5 |
| Neurotoxic | 20 | 0 | 0 | 1,481 | 9 | 12 |
| Hemolytic | 4,279 | 0 | 2 | 9 | 4,867 | 151 |
| Cytolysis | 247 | 0 | 5 | 12 | 151 | 341 |

**Supplementary Table S12**: Directional positive co-annotation percentages across the six non-root MAOMAO toxicity endpoints. Percentages were calculated relative to the total number of positive sequences assigned to each row endpoint and are rounded to two decimal places.

| Row endpoint | Cytotoxic | Embryotoxic | Ichthyotoxic | Neurotoxic | Hemolytic | Cytolysis |
|---|---|---|---|---|---|---|
| Cytotoxic | 100.00% | 0.00% | 0.06% | 0.43% | 91.10% | 5.26% |
| Embryotoxic | 0.00% | 100.00% | 0.00% | 0.00% | 0.00% | 0.00% |
| Ichthyotoxic | 60.00% | 0.00% | 100.00% | 0.00% | 40.00% | 100.00% |
| Neurotoxic | 1.35% | 0.00% | 0.00% | 100.00% | 0.61% | 0.81% |
| Hemolytic | 87.92% | 0.00% | 0.04% | 0.18% | 100.00% | 3.10% |
| Cytolysis | 72.43% | 0.00% | 1.47% | 3.52% | 44.28% | 100.00% |

The largest pairwise overlap was observed between *Cytotoxic* and *Hemolytic*, with 4,279 shared positive sequences. These sequences represented 91.10% of all cytotoxic-positive sequences and 87.92% of all hemolytic-positive sequences. This overlap is consistent with the ontology structure and the positive-only hierarchy rule, through which positive hemolytic evidence can support the broader *Cytotoxic* endpoint.

### S3 Metadata schema and provenance fields

To support transparency, reproducibility, and long-term reuse, the metadata generated during MAOMAO construction were organized into a normalized schema covering source-level, endpoint-level, final dataset-level, hierarchy-level, and audit information. Source-level metadata describe the original resources and their retrieval, licensing, annotation content, and record counts. Endpoint-level metadata document effect-specific filtering and evidence integration. Final dataset-level metadata define the structure and state encoding of the sequence-level pivot, whereas hierarchy- and audit-level records document terminology mappings, propagation rules, ambiguity handling, and hierarchy-derived changes. Missing metadata were preserved explicitly rather than inferred.

#### S3.1 Metadata schema

Source-level metadata keys were normalized to lowercase `snake_case` in the aggregated release metadata. The complete schema preserves the relationships between the original sources, endpoint-specific integration outputs, final sequence–endpoint states, hierarchy rules, ambiguity-support records, and distributed resource files. The principal fields included in the release are summarized in Table **S13**.

**Supplementary Table S13**: Principal metadata fields used for MAOMAO harmonization, provenance tracking, and auditability.

| Field name | Description and allowed values |
|---|---|
| **Source-level metadata** | |
| `source_type` | Type of original resource. Values include Database, Dataset, Publication, or Repository. |
| `resource_behavior` | Indicates whether the resource is static or dynamically updated. |
| `license` | License reported by the original provider; retained as *No information* when unavailable. |
| `publication_year` | Four-digit publication year. |
| `last_update_date` | Last update date reported by the source, when available. |
| `download_date` | Date on which the resource was accessed or downloaded. |
| `file_format` | Format of the retrieved or processed source file, such as CSV, TSV, XLSX, FASTA, JSON, or TXT. |
| `peptide_property` | Peptide activity or toxicity property represented by the source. |
| `annotation_content` | Annotation categories available in the source, such as positive, negative, or unlabeled records. |
| `unit_of_measurement` | Experimental unit reported by the source, when available. |
| `negative_dataset_origin` | Strategy used to define negative examples, such as curated negatives, random selection, or Swiss-Prot sampling. |
| `repository_or_server` | URL of the original repository, server, database, or downloadable resource. |
| `publication` | Literature reference, DOI, PMID, or publication URL associated with the source. |
| `number_of_raw_sequences` | Number of source records before source-specific cleaning or filtering. |
| `number_of_sequences_retained` | Number of records retained after source-specific processing. |
| `number_of_positive_sequences` | Number of retained positive records. |
| `number_of_negative_sequences` | Number of retained negative records. |
| `number_of_unlabel_sequences` | Number of retained records without an explicit positive or negative assignment. |
| `number_of_regression_sequences` | Number of retained quantitative or regression-oriented records, when applicable. |
| `number_of_erroneous_sequences` | Number of sequences removed because of invalid or erroneous content. |
| `number_of_modified_sequences` | Number of modified peptide sequences detected in the source. |
| `number_of_erroneous_modified_ sequences` | Number of modified sequences excluded as erroneous. |
| `modified_sequences_included` | Indicates whether modified peptide sequences were retained. |
| **Endpoint-level metadata** | |
| `task` | Toxicity endpoint represented by the endpoint-integration record. |
| `generated_at` | ISO date and time when the endpoint metadata were generated. |
| `sources.n_unique_sequences` | Number of unique sequences contributed by each source to the endpoint-specific integration. |
| `filters.canonical_residues` | Canonical-residue filter configuration and application status. |
| `filters.length_filter` | Minimum and maximum accepted lengths and filter-application status. |
| `sequence_statistics.canonical_filter` | Sequence counts before and after canonical-residue filtering. |
| `sequence_statistics.length_filter` | Sequence counts before and after length filtering. |
| `sequence_statistics.length_ distribution` | Minimum, maximum, mean, and median peptide lengths. |
| `statistics.total_sequences_final` | Number of final sequences retained for the endpoint before construction of the complete pivot. |
| `statistics.positive` | Summary of positive source evidence and source-label combinations. |
| `statistics.negative` | Summary of negative source evidence and source-label combinations. |
| `statistics.ambiguous` | Number and categories of ambiguous endpoint annotations. |
| `organism_statistics.targets` | Organism names associated with endpoint annotations and their counts, when available. |
| `evidence_negative_dataset_ statistics.category` | Provenance-level characterization of negative evidence, including strong, weak, or unconfirmed negatives. |
| **Final dataset-level metadata** | |
| `dataset.name` | Name of the final sequence-level pivot. |
| `dataset.identifier_column` | Identifier column and sequential identifier format. |
| `dataset.sequence_column` | Name of the peptide-sequence column. |

*Continued on next page*

**Supplementary Table S13**: Principal metadata fields used for MAOMAO harmonization, provenance tracking, and auditability. Continued.

| Field name | Description and allowed values |
|---|---|
| `dataset.annotation_columns` | Seven controlled endpoint columns included in the pivot. |
| `dataset.n_unique_sequences` | Number of unique peptide sequences: 71,701. |
| `dataset.n_sequence_endpoint_rows` | Size of the complete sequence–endpoint cross-product: 501,907 combinations. |
| `dataset.label_encoding` | State encoding: 1 = positive, 0 = negative, 2 = ambiguous, 3 = unlabeled, and 999 = no information. |
| `dataset.state_priority` | States that are ambiguous for a specific endpoint cannot be overwritten by hierarchical positivity. |
| `dataset.negative_evidence_policy` | Negative evidence is endpoint-specific and is not propagated through the hierarchy. |
| `dataset.missing_information_policy` | Code 999 denotes unavailable endpoint information and is distinct from negative and unlabeled evidence. |
| **Hierarchy- and audit-level metadata** | |
| `controlled_vocabulary_and_hierarchy.version` | Version of the MAOMAO controlled vocabulary and hierarchy. |
| `hierarchy_policy.positive_propagation` | Indicates that positive evidence can propagate from a child endpoint to its parent. |
| `hierarchy_policy.negative_propagation` | Indicates that negative evidence is not propagated. |
| `hierarchy_policy.ambiguity_overrides_hierarchy` | Indicates that direct parent ambiguity blocks hierarchy-derived positivity. |
| `terms.term` | Harmonized or source-specific toxicity term. |
| `terms.label_type` | Role of the term, such as broad, intermediate, specific, or source-specific. |
| `terms.parent_terms` | Parent endpoint or endpoints associated with a controlled term. |
| `terms.harmonized_to` | Controlled endpoint to which a source-specific synonym was mapped. |
| `rules_applied.rule_id` | Identifier of the applied terminology or hierarchy rule. |
| `rules_applied.source_terms` | Child or source terms that trigger the rule. |
| `rules_applied.target_term` | Parent or harmonized endpoint modified by the rule. |
| `rules_applied.condition` | Logical condition under which the rule is applied. |
| `rules_applied.action` | State change or terminology mapping performed by the rule. |
| `rules_applied.blocked_when` | Endpoint-specific state that prevents propagation, when applicable. |
| `rules_applied.propagation_policy` | Propagation policy; the hierarchy rules use positive-only propagation. |
| `ambiguity.support_file` | File containing endpoint-specific support summaries for ambiguous sequence–endpoint states. |
| `output_files` | File-level inventory containing descriptions, dimensions, checksums, and column information for principal outputs. |

#### S3.2 Source metadata coverage

Metadata availability was evaluated across the 54 sources integrated into MAOMAO. As summarized in Table **S14**, publication references, repository or server information, download dates, file formats, source types, and resource behavior were available for all sources. Annotation content and last-update dates were available for 53 sources (98.15%). In contrast, explicit licensing information was available for 13 sources (24.07%), and experimental units of measurement were documented for only 2 sources (3.70%). Missing values were retained as unavailable information rather than being inferred from related publications or external resources.

**Supplementary Table S14**: Metadata-field availability across the 54 sources integrated into MAOMAO.

| Metadata field | Available | Total | Coverage |
|---|---|---|---|
| Publication | 54 | 54 | 100% |
| Repository or server | 54 | 54 | 100% |
| Download date | 54 | 54 | 100% |
| File format | 54 | 54 | 100% |
| Source type | 54 | 54 | 100% |
| Resource behavior | 54 | 54 | 100% |
| Annotation content | 53 | 54 | 98.15% |
| Last-update date | 53 | 54 | 98.15% |
| License | 13 | 54 | 24.07% |
| Unit of measurement | 2 | 54 | 3.70% |

The metadata-coverage results reflect the information explicitly reported by the original resources and available at the time of retrieval. Consequently, low coverage for fields such as licensing and measurement units represents limitations of the source documentation rather than values omitted during MAOMAO harmonization.

#### S3.3 Source access and licensing audit

Access and licensing information was consolidated for the 54 resources integrated into MAOMAO (Table **S15**). An explicit license was recorded for 13 resources (24.07%), whereas no explicit license was identified in the recorded metadata for the remaining 41 resources.

The table summarizes the recorded retrieval method, last-update information, reported license or terms, and a provisional assessment of redistribution and attribution requirements. Interpretations of the MIT, Apache License 2.0, Creative Commons Attribution 4.0, GNU General Public License, and Database Contents License/Open Database License conditions were based on their respective official license texts [Open Source Initiative, 2026, Apache Software Foundation, 2004, Creative Commons, 2013, Free Software Foundation, 2007, Open Data Commons, 2008, 2009]. Because a license displayed at the repository level may apply to software or documentation rather than to the associated data files, all redistribution assessments should be interpreted as provisional and subject to source-level verification.

**Supplementary Table S15**: Source retrieval and provisional licensing audit for the 54 resources integrated into MAOMAO. Reported licenses reproduce the information recorded in the source metadata.

| Source | Last update | Retrieval method | Reported license or terms | Redistribution status | Attribution or conditions |
|---|---|---|---|---|---|
| AMPDB v1 | 2023-05-03–2023-05-26 | Database/web-server download | No information | Not established | Not established |
| AMPDeep | 2022-09-28 | Repository download | MIT | Permitted, subject to license terms | Copyright and permission notice |
| Abdelbaky et al. | 2024-06-29 | Repository download | No information | Not established | Not established |
| Almotairi et al. | 2024-06-28 | Repository download | No information | Not established | Not established |
| BIOPEP-UWM | 2025-08-27 | Database/web-server download | No information | Not established | Not established |
| Bhatnagar et al. | 2024-10-30 | Supplementary-file retrieval | No information | Not established | Not established |
| BiToxNet | 2025-11-09 | Repository download | No information | Not established | Not established |
| CAPTP | 2025-09-24 | Repository download | No information | Not established | Not established |
| CICERON | 2024-04-30 | Repository download | No information | Not established | Not established |
| ConsAMPHemo | 2024-09-28 | Repository download | No information | Not established | Not established |
| DRAMP | 2025-07-16 | Database/web-server download | Creative Commons Attribution 4.0 | Permitted with attribution | Credit, license reference, and change indication |
| HAPPENN | No information | Database/web-server download | No information | Not established | Not established |
| HEPAD | 2024-04-21 | Repository download | No information | Not established | Not established |
| HLPpred-Fuse | 2020-04-20 | Database/web-server download | No information | Not established | Not established |
| HMAMP-main | 2025-09-03 | Repository download | No information | Not established | Not established |
| HemoDL | 2023-04-18 | Repository download | No information | Not established | Not established |
| HemoFuse | 2024-09-19 | Repository download | No information | Not established | Not established |
| HemoPI | 2016-03-08 | Database/web-server download | No information | Not established | Not established |
| HemoPI2.0 | 2025-02-05 | Database/web-server download | No information | Not established | Not established |
| Hemolytic-Pred | 2023-07-05 | Supplementary-file retrieval | No information | Not established | Not established |
| Hemolytik | 2014-01-01 | Database/web-server download | No information | Not established | Not established |
| HyPepTox-Fuse | 2025-07-24 | Repository download | Apache 2.0 | Permitted, subject to license terms | License and applicable NOTICE retention |
| Karasev et al. | 2024-12-01 | Supplementary-file retrieval | No information | Not established | Not established |
| MultiPep | 2023-03-30 | Repository download | MIT | Permitted, subject to license terms | Copyright and permission notice |
| Multitox | 2025-05-19 | Repository download | No information | Not established | Not established |
| NTXpred | 2007-11-01 | Database/web-server download | No information | Not established | Not established |
| NTXpred2 | 2025-02-21 | Database/web-server download | GNU general public license | Permitted under GPL conditions | License notices and applicable copyleft/source obligations |
| PLPTP | 2025-02-23 | Repository download | No information | Not established | Not established |
| Pep-Lab_db | 2023-01-10 | Database/web-server download | No information | Not established | Not established |
| PeptiTox | 2025-03-28 | Repository download | No information | Not established | Not established |
| Peptipedia2.0 | 2024-03-23 | Database/web-server download | DbCL | Conditional under DbCL/ODbL terms | Applicable attribution and database-license conditions |
| Plisson et al. | 2020-08-24 | Repository download | MIT | Permitted, subject to license terms | Copyright and permission notice |
| ProToxin | 2025-06-25 | Database/web-server download | No information | Not established | Not established |
| QSVM-PEPTIDE | 2024-09-19 | Repository download | No information | Not established | Not established |
| SATPdb | 2015-11-01 | Database/web-server download | No information | Not established | Not established |
| StrucToxNet | 2025-06-05 | Repository download | No information | Not established | Not established |
| TPpred-LE | 2023-01-31 | Database/web-server download | No information | Not established | Not established |

*Continued on next page*

**Supplementary Table S15** – *Continued from previous page*

| Source | Last update | Retrieval method | Reported license or terms | Redistribution status | Attribution or conditions |
|---|---|---|---|---|---|
| ToxDL 2.0 | 2025-04-02 | Database/web-server download | No information | Not established | Not established |
| ToxGIN | 2025-01-18 | Repository download | No information | Not established | Not established |
| ToxIBTL | 2022-01-24 | Repository download | No information | Not established | Not established |
| ToxMSRC | 2025-08-25 | Repository download | No information | Not established | Not established |
| ToxTeller | 2023-12-25 | Repository download | No information | Not established | Not established |
| ToxiPep | 2025-05-20 | Repository download | No information | Not established | Not established |
| ToxinPred | 2013-09-13 | Database/web-server download | No information | Not established | Not established |
| ToxinPred 2.0 | 2022-05-21 | Database/web-server download | GNU general public license | Permitted under GPL conditions | License notices and applicable copyleft/source obligations |
| ToxinPred 3.0 | 2024-07-21 | Database/web-server download | GNU general public license | Permitted under GPL conditions | License notices and applicable copyleft/source obligations |
| UniDL4BioPep | 2024-11-22 | Repository download | MIT | Permitted, subject to license terms | Copyright and permission notice |
| Zhao et al. | 2021-05-26 | Supplementary-file retrieval | Creative Commons Attribution 4.0 | Permitted with attribution | Credit, license reference, and change indication |
| csm-toxin | 2023-01-30 | Repository download | No information | Not established | Not established |
| hemolytik 2.0 | 2014-05-14 | Database/web-server download | No information | Not established | Not established |
| hemonet | 2021-03-11 | Repository download | No information | Not established | Not established |
| iAMPCN | 2023-03-02 | Repository download | No information | Not established | Not established |
| peptidereactor | 2021-05-25 | Repository download | MIT | Permitted, subject to license terms | Copyright and permission notice |
| tAMPer | 2023-11-19 | Repository download | GNU general public license | Permitted under GPL conditions | License notices and applicable copyleft/source obligations |

*Note.* "No information" indicates that an explicit license was not recorded in the source metadata and must not be interpreted as public-domain status or permission to redistribute. The redistribution and attribution columns constitute a provisional assessment based on the reported license label and require source-level verification that the stated license applies to the downloaded data rather than only to repository software or documentation. Recorded retrieval methods correspond to repository downloads, database or web-server downloads, and supplementary-file retrieval. Associated publications are provided in the reference list, whereas source locations are preserved in the released provenance metadata.

### S4 Computational resource generation and file organization

#### S4.1 Descriptor Layer generation

A single descriptor table was generated from the final 71,701-sequence MAOMAO pivot to provide reusable physicochemical and sequence-derived features across all toxicity endpoints. Descriptor extraction was performed with the Roxy framework v0.1.0 [KrenAI-Lab, 2026b] using the peptide sequence column as input. No single target column was specified because the resource contains seven endpoint-state columns. AAindex features were not enabled; the descriptor configuration retained the sequence-global feature set.

The output table contains the sequence identifier, peptide sequence, seven endpoint-state columns, and 41 descriptor columns, resulting in 71,701 rows and 50 columns. The descriptor matrix is distributed as `dataset_characterization/sequence_descriptors.csv`, together with `dataset_characterization/metadata.json`, which documents the generation settings, state encoding, output dimensions, descriptor names, and quality-control results.

**Supplementary Table S16**: Descriptor Layer generation settings.

| Parameter | Value |
|---|---|
| Descriptor framework | Roxy v0.1.0 |
| Input resource | Final MAOMAO sequence-level pivot |
| Input rows | 71,701 unique peptide sequences |
| Identifier column | `id` |
| Sequence column | `sequence` |
| Endpoint-state columns | `toxic`, `cytotoxic`, `hemolytic`, `cytolysis`, `neurotoxic`, `embryotoxic`, and `ichthyotoxic` |
| Feature set | Sequence-global descriptors |
| AAindex features | Disabled |
| Target column | None; all seven endpoint-state columns were retained |
| Descriptor columns | 41 |
| Output dimensions | 71,701 rows × 50 columns |
| Descriptor output | `dataset_characterization/sequence_descriptors.csv` |

*Continued on next page*

**Supplementary Table S16** – *Continued from previous page*

| Parameter | Value |
|---|---|
| Metadata output | `dataset_characterization/metadata.json` |

The 41 descriptors are organized into the groups summarized in Table **S17**.

**Supplementary Table S17**: Sequence-derived descriptors included in the MAOMAO Descriptor Layer.

| Descriptor group | Descriptor columns | Interpretation |
|---|---|---|
| Sequence length | `length` | Number of amino-acid residues in the peptide. |
| Amino-acid composition | `aa_frac_A`, `aa_frac_C`, `aa_frac_D`, `aa_frac_E`, `aa_frac_F`, `aa_frac_G`, `aa_frac_H`, `aa_frac_I`, `aa_frac_K`, `aa_frac_L`, `aa_frac_M`, `aa_frac_N`, `aa_frac_P`, `aa_frac_Q`, `aa_frac_R`, `aa_frac_S`, `aa_frac_T`, `aa_frac_V`, `aa_frac_W`, `aa_frac_Y` | Fraction of each canonical amino acid in the sequence. |
| Residue-class composition | `frac_aromatic`, `frac_positive`, `frac_negative`, `frac_polar`, `frac_nonpolar` | Fractions of residues grouped by physicochemical class. |
| Hydrophobicity and hydropathy | `gravy_kd`, `hydropathy_eisenberg` | Mean sequence hydropathy according to the Kyte–Doolittle and Eisenberg scales. |
| Structural propensity and disorder | `top_idp_mean`, `helix_propensity_mean`, `sheet_propensity_mean` | Mean disorder, helix, and sheet propensity values. |
| Interaction propensity | `boman_index` | Estimated protein-binding or interaction potential. |
| Charge-related properties | `net_charge_pH`, `fcr`, `ncpr` | Estimated net charge, fraction of charged residues, and net charge per residue. |
| Hydrogen-bonding capacity | `donors_per_residue`, `acceptors_per_residue` | Estimated hydrogen-bond donors and acceptors normalized by sequence length. |
| Sequence complexity | `aa_entropy`, `lc_k1`, `lc_k2`, `lc_k3` | Amino-acid entropy and low-complexity measures at increasing word lengths. |

#### S4.2 Numerical representation extraction

Numerical representations were generated once for the complete harmonized sequence collection and were subsequently reused across endpoint-specific benchmark construction. Protein language model embeddings were obtained with the Sylphy v0.2.0 framework [KrenAI-Lab, 2026c] using pretrained models without task-specific fine-tuning. Residue-level outputs from the final embedding layer were mean-pooled to obtain one fixed-length vector per peptide sequence.

Embedding extraction used GPU acceleration with `cuda`, `fp32` precision, and a batch size of 16. For each model, `embeddings.csv` stores the numerical matrix and `full_data.csv` preserves the correspondence between identifiers, peptide sequences, endpoint states, and embedding rows. The representation root is `numerical_representation_data/maomao/`.

**Supplementary Table S18**: Embedding extraction settings used for the protein language model representations.

| Parameter | Value |
|---|---|
| Framework | Sylphy v0.2.0 |
| Input resource | Complete MAOMAO sequence collection |
| Model state | Pretrained, without task-specific fine-tuning |
| Representation layer | Final embedding layer |
| Pooling strategy | Mean pooling across residue embeddings |
| Device | `cuda` |
| Precision | `fp32` |
| Batch size | 16 |
| Output format | CSV |
| Main output file | `embeddings.csv` |
| Companion file | `full_data.csv` |

#### S4.3 Supported protein language model embeddings

The MAOMAO Embedding Layer provides ten precomputed sequence-level embedding matrices derived from pretrained protein language models. These representations cover five model families—Ankh, ESM2, ESM-C, Mistral-Prot, and ProtTrans—and include architectures with different parameter scales and output dimensions. Each matrix assigns a fixed-length numerical representation to the 71,701 peptide sequences in the final MAOMAO pivot, allowing the same curated sequence collection to be examined across alternative learned representation spaces without requiring users to regenerate the embeddings.

Table **S19** summarizes the model variant, protein language model family, relative file location, and embedding dimension of each distributed matrix. The available representations range from 320 to 1,536 features per peptide sequence. The four ESM2 variants provide representations obtained from models of increasing scale, whereas the Ankh, ESM-C, Mistral-Prot, and ProtTrans models broaden the representation layer across distinct pretrained protein language model families.

**Supplementary Table S19**: Protein language model embeddings included in the MAOMAO Embedding Layer. File paths are relative to `numerical_representation_data/maomao/sylphy_embedding/`.

| Model | Family | Embedding file | Dimension |
|---|---|---|---|
| Ankh2-ext1 [Elnaggar et al., 2023] | Ankh | `ankh2_ext1/embeddings.csv` | 1536 |
| Ankh3-large [Alsamkary et al., 2025] | Ankh | `ankh3_large/embeddings.csv` | 1536 |
| ESM2 t6 8M UR50D [Lin et al., 2022] | ESM2 | `esm2_t6_8M_UR50D/embeddings.csv` | 320 |
| ESM2 t12 35M UR50D [Lin et al., 2022] | ESM2 | `esm2_t12_35M_UR50D/embeddings.csv` | 480 |
| ESM2 t30 150M UR50D [Lin et al., 2022] | ESM2 | `esm2_t30_150M_UR50D/embeddings.csv` | 640 |
| ESM2 t33 650M UR50D [Lin et al., 2022] | ESM2 | `esm2_t33_650M_UR50D/embeddings.csv` | 1280 |
| ESM-C 300M [Candido et al., 2026] | ESM-C | `esmc_300m/embeddings.csv` | 960 |
| Mistral-Prot v1 134M [Mourad, 2024] | Mistral-Prot | `mistral_prot_v1_134M/embeddings.csv` | 768 |
| ProtBERT [Elnaggar et al., 2021] | ProtTrans | `prot_bert/embeddings.csv` | 1024 |
| ProtT5-XL-UniRef50 [Elnaggar et al., 2021] | ProtTrans | `prot_t5_xl_uniref50/embeddings.csv` | 1024 |

##### S4.4 One-hot baseline representation

A conventional one-hot representation was generated using Sylphy v0.2.0 as a baseline and was distributed separately from the protein language model embeddings. In this encoding, each amino acid at a given sequence position is represented by a binary indicator vector over the amino acid alphabet, providing a direct sequence-based representation without learned contextual information [Jing et al., 2019]. The encoded matrix is stored under `numerical_representation_data/maomao/sylphy_one_hot/one_hot/encoded.csv`, whereas `full_data.csv` retains the corresponding sequence identifiers, peptide sequences, endpoint states, and row correspondence. Together, the ten protein language model embedding matrices and the one-hot baseline define the eleven numerical representations available for reconstructing representation-specific benchmark datasets.

##### S4.5 Benchmark partition generation

Benchmark-ready partitions were generated with BioSieve v0.1.0 [KrenAI-Lab, 2026a] for toxicity endpoints containing sufficient positive and negative sequence counts to support binary five-fold partitioning. Eligibility was assessed using the final endpoint states after hierarchy application. Ambiguous, unlabeled, and no-information states were not treated as binary benchmark classes. As summarized in Table **S20**, four endpoints met the predefined eligibility criteria: *Toxic*, *Cytotoxic*, *Hemolytic*, and *Neurotoxic*. *Cytolysis*, *Embryotoxic*, and *Ichthyotoxic* were not included because their final positive–negative distributions were insufficient for the predefined five-fold design.

**Supplementary Table S20**: Endpoint eligibility for binary benchmark partitioning. Positive and negative counts correspond to the final pivot after hierarchy application.

| Endpoint | Positive | Negative | Eligible | Benchmark interpretation |
|---|---|---|---|---|
| *Toxic* | 12,049 | 37,129 | Yes | Sufficient positive and negative examples for the predefined design. |
| *Cytotoxic* | 4,697 | 19,662 | Yes | Sufficient positive and negative examples for the predefined design. |
| *Hemolytic* | 4,867 | 19,350 | Yes | Sufficient positive and negative examples for the predefined design. |
| *Neurotoxic* | 1,481 | 1,700 | Yes | Sufficient positive and negative examples for the predefined design. |
| *Cytolysis* | 341 | 2 | No | The negative class is too small for reliable five-fold partitioning. |
| *Embryotoxic* | 2 | 0 | No | No negative endpoint-specific examples are available. |
| *Ichthyotoxic* | 5 | 0 | No | No negative endpoint-specific examples are available. |

For each eligible endpoint, random and stratified partitions were generated once for each of 30 random seeds. Because these partitioning strategies do not depend on numerical features, the resulting train, validation, and test assignments were stored independently of the 11 numerical representations. Random splitting did not explicitly preserve endpoint-label proportions, whereas stratified splitting preserved class proportions across the train, validation, and test subsets. The same set of seeds was used across endpoints and partitioning strategies to support paired comparisons. Representation-specific benchmark datasets can be reconstructed by joining each partition file with the selected numerical representation using the sequence identifier. The complete partition-generation settings are summarized in Table **S21**.

**Supplementary Table S21**: Benchmark partition generation settings used in the MAOMAO release.

| **Parameter** | **Value** |
|---|---|
| Partitioning framework | BioSieve v0.1.0 |
| Eligible endpoints | *Toxic*, *Cytotoxic*, *Hemolytic*, and *Neurotoxic* |
| Endpoint classes | Positive and negative |
| Compatible numerical representations | Ten protein language model embeddings and one one-hot baseline |
| Representation count | 11 |
| Split source | Non-reduced endpoint-specific binary datasets: `no_reduced` |
| Enabled strategies | `random_kfold` and `stratified_kfold` |
| Number of folds | 5 |
| Random seeds | 30 |
| Seed file | `general_configs/random_seeds_30.csv` |
| Shuffle | Enabled |
| Validation size | 0.1 |
| Stratification variable | Endpoint label |
| Required fold files | `train.csv`, `val.csv`, and `test.csv` |
| Stored fold columns | `id` and `label` |
| Validation criteria | At least two classes overall and at least two classes in every generated split |
| Invalid split handling | Invalid configurations are documented without stopping the complete workflow |
| Stored partition configuration count | 240 |
| Stored fold count | 1,200 |
| Reconstructable representation-specific configuration count | 2,640 |
| Reconstructable representation-specific fold count | 13,200 |

The number of physically stored partition configurations was calculated according to Equation S1:

$$N_{\text{stored}} = 4 \text{ endpoints} \times 30 \text{ seeds} \times 2 \text{ strategies} = 240. \tag{S1}$$

Because each stored configuration contains five folds, the Benchmark Layer contains:

$$N_{\text{stored folds}} = 240 \text{ configurations} \times 5 \text{ folds} = 1{,}200. \tag{S2}$$

Each stored partition can be combined with any of the 11 numerical representations. The number of reconstructable representation-specific benchmark configurations is therefore:

$$N_{\text{reconstructed}} = 240 \text{ configurations} \times 11 \text{ representations} = 2{,}640. \tag{S3}$$

The corresponding number of representation-specific train–validation–test fold instances is:

$$N_{\text{reconstructed folds}} = 1{,}200 \text{ folds} \times 11 \text{ representations} = 13{,}200. \tag{S4}$$

#### S4.6 File organization

The MAOMAO release is organized into three principal groups of computational artifacts: dataset characterization files, numerical sequence representations, and benchmark-ready data partitions. This organization separates the description of the curated biological resource from the numerical encodings and partition files generated for downstream machine-learning experiments. Directory and file names are preserved across the release to maintain correspondence between sequences, endpoint annotations, representations, and benchmark configurations.

The `dataset_characterization/` directory contains the resource-level metadata and the calculated sequence descriptors. The `metadata.json` file documents the dataset configuration, processing parameters, provenance information, and summary statistics associated with the released resource. The `sequence_descriptors.csv` file contains the sequence identifiers, peptide sequences, endpoint states, and calculated physicochemical and sequence-derived descriptors.

```
dataset_characterization/
  metadata.json
  sequence_descriptors.csv
```

Precomputed numerical representations are distributed under `numerical_representation_data/maomao/`. Protein language model embeddings and the one-hot baseline are stored in separate branches. For each protein language model, `embeddings.csv` contains the fixed-length sequence-level representation matrix, whereas `full_data.csv` preserves the associated sequence identifiers, peptide sequences, endpoint states, and row correspondence. The one-hot branch follows the same principle: `encoded.csv` contains the numerical encoding and `full_data.csv` contains the corresponding sequence-level information.

**Supplementary Table S22** – *Continued from previous page*

| File | Rows | Columns | Description |
|---|---|---|---|
| audit_endpoint_counts.csv | 7 | 5 | Positive, negative, ambiguous, and unlabeled counts for each endpoint. |
| audit_hierarchy_changes.csv | 11,831 | 9 | Sequence-level audit of hierarchy support, inference, ambiguity blocking, and conflicts. |
| metadata.json | — | — | Aggregated resource, source, endpoint, hierarchy, ambiguity, sequence, and file metadata. |

#### S5.2 Descriptor Layer outputs

The Descriptor Layer contains one 71,701-row table combining the sequence identifiers, peptide sequences, seven endpoint-state columns, and 41 numerical descriptors. The accompanying metadata file records descriptor definitions, generation parameters, dimensions, label encoding, and quality-control results.

**Supplementary Table S23**: Descriptor Layer file summary.

| Resource component | File | Description |
|---|---|---|
| Descriptor matrix | dataset_characterization/sequence_descriptors.csv | 71,701 rows and 50 columns, including 41 descriptors. |
| Descriptor metadata | dataset_characterization/metadata.json | Generation settings, descriptor names, state distributions, and quality-control information. |

#### S5.3 Embedding Layer outputs

The Embedding Layer contains ten protein language model-derived matrices and one one-hot baseline representation. Protein language model files are stored under model-specific folders, and each matrix is accompanied by a file preserving identifier and sequence correspondence.

**Supplementary Table S24**: Numerical representation file summary.

| Resource component | Directory or file pattern | Summary |
|---|---|---|
| Protein language model embeddings | numerical_representation_data/maomao/sylphy_embedding/<model>/ | Ten model-specific embedding matrices. |
| Main embedding file | embeddings.csv | One fixed-length vector per peptide sequence. |
| Embedding companion file | full_data.csv | Identifier, sequence, endpoint-state, and row correspondence. |
| One-hot baseline | numerical_representation_data/maomao/sylphy_one_hot/one_hot/encoded.csv | Conventional sequence encoding used as the eleventh benchmark representation. |

#### S5.4 Benchmark partition resources

The Benchmark Layer contains 240 stored endpoint–seed–strategy partition configurations and 1,200 stored train–validation–test folds. Each stored configuration contains fifteen fold-level CSV files and six seed-level configuration, report, completion, and execution-log files, corresponding to 5,040 benchmark-layer files under the released directory schema. The fold-level files contain only sequence identifiers and binary labels. By combining the stored partitions with the 11 numerical representations distributed in the Embedding Layer, users can reconstruct 2,640 representation-specific benchmark configurations and 13,200 representation-specific fold instances.

**Supplementary Table S25**: Benchmark partition resource summary.

| Benchmark component | Summary |
|---|---|
| Benchmark directory | split_process/maomao_<endpoint>/no_reduced/ |
| Eligible toxicity endpoints | Four: *Toxic*, *Cytotoxic*, *Hemolytic*, and *Neurotoxic* |
| Compatible numerical representations | Eleven: ten protein language model embeddings and one one-hot baseline |
| Random seeds | 30 |
| Partitioning strategies | random_kfold and stratified_kfold |
| Stored partition configuration count | $4 \times 30 \times 2 = 240$ |
| Fold configuration | Five train–validation–test folds per stored configuration |
| Stored fold count | $240 \times 5 = 1,200$ |
| Reconstructable representation-specific configuration count | $240 \times 11 = 2,640$ |

*Continued on next page*

**Supplementary Table S25** – *Continued from previous page*

| **Benchmark component** | **Summary** |
|---|---|
| Reconstructable representation-specific fold count | $1,200 \times 11 = 13,200$ |
| Fold-level files | `train.csv`, `val.csv`, and `test.csv` |
| Fold-level columns | `id` and `label` |
| Seed-level files | Logs, completion marker, parameters, split report, and summary |
| Benchmark-layer file count | 5,040 files under the stored partition schema |

#### S5.5 Release-level organization

The MAOMAO release is organized into complementary resource layers according to the function of each component within the data-construction and reuse workflow. This modular organization separates the harmonized biological data from derived numerical representations, benchmark resources, workflow implementations, configuration files, and supporting documentation, while preserving consistent identifiers and directory relationships across layers. As summarized in Table **S26**, each layer is associated with a defined set of directories and files that supports data traceability, independent reuse, and reproducible regeneration of the resource components.

**Supplementary Table S26**: Release-level organization of MAOMAO resource components.

| **Resource layer** | **Directory** | **Main content** |
|---|---|---|
| Core Layer | `processed_data/`; source and endpoint metadata directories | Final pivot, source-processed datasets, endpoint integration outputs, ambiguity support, hierarchy audits, and metadata. |
| Descriptor Layer | `dataset_characterization/` | Global sequence-descriptor table and descriptor metadata. |
| Embedding Layer | `numerical_representation_data/` | Ten protein language model embeddings and one one-hot baseline. |
| Benchmark Layer | `split_process/` | Endpoint-specific, representation-independent random and stratified five-fold partitions, compact identifier–label files, reports, parameters, summaries, and logs. |
| Configuration files | `general_configs/` | Random seed list and shared configuration resources. |
| Workflow resources | `notebooks_and_scripts/`; `pipelines/`; `src/` | Data acquisition, parsing, harmonization, characterization, representation generation, and partitioning workflows. |
| Documentation Layer | Root-level and documentation files | README files, controlled vocabulary descriptions, licensing information, versioning records, and resource inventories. |

#### S5.6 Release, versioning, and licensing

The MAOMAO resource is distributed through a versioned Zenodo record, with each release preserving the harmonized data, derived numerical representations, benchmark resources, metadata, and supporting documentation. MAOMAO-derived data products are distributed under the Creative Commons Attribution 4.0 International license. Original third-party source files remain subject to the licensing terms of their respective providers. Source code and reproducible workflows are distributed separately through the project GitHub repository under the MIT License.

Hazem Alsamkary, Mohamed Elshaffei, Mohamed Elkerdawy, and Ahmed Elnaggar. Ankh3: Multi-task pretraining with sequence denoising and completion enhances protein representations. *arXiv preprint arXiv:2505.20052*, 2025.

Mehrad Ansari and Andrew D White. Learning peptide properties with positive examples only. *bioRxiv*, pages 2023–06, 2023a.

Mehrad Ansari and Andrew D White. Serverless prediction of peptide properties with recurrent neural networks. *Journal of Chemical Information and Modeling*, 63(8):2546–2553, 2023b.

Apache Software Foundation. Apache license, version 2.0, 2004. URL `https://www.apache.org/licenses/LICENSE-2.0`. Consultado: 27 de julio de 2026.

Özlem Sultan Aslantürk. In vitro cytotoxicity and cell viability assays: principles, advantages, and disadvantages. In *Genotoxicity-A predictable risk to our actual world*. IntechOpen, 2017.

Nancy Baker, Alan Boobis, Lyle Burgoon, Edward Carney, Richard Currie, Ellen Fritsche, Thomas Knudsen, Madeleine Laffont, Aldert H Piersma, Alan Poole, et al. Building a developmental toxicity ontology. *Birth defects research*, 110(6):502–518, 2018.

Kathryn Bambino and Jaime Chu. Zebrafish in toxicology and environmental health. *Current topics in developmental biology*, 124: 331–367, 2017.

Pranshul Bhatnagar, Yashi Khandelwal, Shagun Mishra, Arnab Dutta, Debirupa Mitra, Swati Biswas, et al. Predicting antibacterial activity, efficacy, and hemotoxicity of peptides using an explainable machine learning framework. *Process Biochemistry*, 145:163–174, 2024.

Edoardo Bizzotto, Guido Zampieri, Laura Treu, Pasquale Filannino, Raffaella Di Cagno, and Stefano Campanaro. Classification of bioactive peptides: a systematic benchmark of models and encodings. *Computational and Structural Biotechnology Journal*, 23: 2442–2452, 2024.

Gabriel Cabas-Mora, Anamaría Daza, Nicole Soto-García, Valentina Garrido, Diego Alvarez, Marcelo Navarrete, Lindybeth Sarmiento-Varón, Julieta H Sepúlveda Yañez, Mehdi D Davari, Frederic Cadet, et al. Peptipedia v2. 0: a peptide sequence database and user-friendly web platform. a major update. *Database*, 2024:baae113, 2024.

Salvatore Candido, Thomas Hayes, Alexander Derry, Roshan Rao, Zeming Lin, Robert Verkuil, Bryan Wu, Jin Sub Lee, Elise S. Bruguera, Jehan A. Keval, Mykhailo Kopylov, John E. Pak, Wesley Wu, Neil Thomas, Samson Mataraso, Alvin Hsu, Ashton C. Trotman-Grant, Kilian Fatras, Allan dos Santos Costa, Rohil Badkundri, Halil Akın, Deniz Oktay, Jonathan Deaton, Elizabeth Montabana, Hrishita Sitwala, Yue Yu, Marius Wiggert, Dylan Alexander Carlin, Anthony W. Goering, Tomasz Blazejewski, McCullen Sandora, Michael Hla, Tina Z. Jia, Leon H. Kloker, Nicholas J. Sofroniew, Masatoshi Uehara, Jassi Pannu, Sharrol Bachas, Daniel S. Liu, Tom Sercu, and Alexander Rives. Language modeling materializes a world model of protein biology, 2026. URL `https://www.biorxiv.org/content/10.64898/2026.06.03.729735`. Preprint.

Kevin Castillo-Mendieta, Guillermin Aguero-Chapin, Edgar Marquez, Yunierkis Perez-Castillo, Stephen J Barigye, Mariela Pérez-Cárdenas, Facundo Peréz-Giménez, and Yovani Marrero-Ponce. Multiquery similarity searching models: an alternative approach for predicting hemolytic activity from peptide sequence. *Chemical Research in Toxicology*, 37(4):580–589, 2024a.

Kevin Castillo-Mendieta, Guillermin Agüero-Chapin, Edgar A Marquez, Yunierkis Perez-Castillo, Stephen J Barigye, Nelson Santiago Vispo, Cesar R García-Jacas, and Yovani Marrero-Ponce. Peptide hemolytic activity analysis using visual data mining of similarity-based complex networks. *npj Systems Biology and Applications*, 10(1):115, 2024b.

Kumardeep Chaudhary, Ritesh Kumar, Sandeep Singh, Abhishek Tuknait, Ankur Gautam, Deepika Mathur, Priya Anand, Grish C Varshney, and Gajendra PS Raghava. A web server and mobile app for computing hemolytic potency of peptides. *Scientific reports*, 6(1):22843, 2016.

Sih-Han Chen, Jen-Chieh Yu, Yi-Hsiang Lin, Shao-Chun Kuo, Kuan Ni, and Ching-Tai Chen. Hepad: enhancing hemolytic peptide prediction with adaptive feature engineering and diverse sequence descriptors. *BMC bioinformatics*, 26(1):234, 2025.

Creative Commons. Creative commons attribution 4.0 international public license, 2013. URL `https://creativecommons.org/licenses/by/4.0/legalcode`. Consultado: 27 de julio de 2026.

Durdam Das, Mohini Jaiswal, Fatima Nazish Khan, Shahzaib Ahamad, and Shailesh Kumar. Plantpepdb: A manually curated plant peptide database. *Scientific reports*, 10(1):2194, 2020.

Zhenjiao Du, Xingjian Ding, Yixiang Xu, and Yonghui Li. Unidl4biopep: a universal deep learning architecture for binary classification in peptide bioactivity. *Briefings in Bioinformatics*, 24(3):bbad135, 2023.

Hossein Ebrahimikondori, Darcy Sutherland, Anat Yanai, Amelia Richter, Ali Salehi, Chenkai Li, Lauren Coombe, Monica Kotkoff, René L Warren, and Inanc Birol. Structure-aware deep learning model for peptide toxicity prediction. *Protein Science*, 33(7):e5076, 2024.

Ahmed Elnaggar, Michael Heinzinger, Christian Dallago, Ghalia Rehawi, Yu Wang, Llion Jones, Tom Gibbs, Tamas Feher, Christoph Angerer, Martin Steinegger, et al. Prottrans: toward understanding the language of life through self-supervised learning. *IEEE transactions on pattern analysis and machine intelligence*, 44(10):7112–7127, 2021.

Ahmed Elnaggar, Hazem Essam, Wafaa Salah-Eldin, Walid Moustafa, Mohamed Elkerdawy, Charlotte Rochereau, and Burkhard Rost. Ankh: Optimized protein language model unlocks general-purpose modelling. *arXiv preprint arXiv:2301.06568*, 2023.

Free Software Foundation. GNU general public license, version 3, 2007. URL `https://www.gnu.org/licenses/gpl-3.0.html`. Consultado: 27 de julio de 2026.

Shun Gao, Yanna Jia, Feifei Cui, Junlin Xu, Yajie Meng, Leyi Wei, Qingchen Zhang, Quan Zou, and Zilong Zhang. Plptp: a motif-based interpretable deep learning framework based on protein language models for peptide toxicity prediction. *Journal of Molecular Biology*, 437(12):169115, 2025.

Ankur Gautam, Kumardeep Chaudhary, Sandeep Singh, Anshika Joshi, Priya Anand, Abhishek Tuknait, Deepika Mathur, Grish C Varshney, and Gajendra PS Raghava. Hemolytik: a database of experimentally determined hemolytic and non-hemolytic peptides. *Nucleic acids research*, 42(D1):D444–D449, 2014.

Alexander GB Grønning, Tim Kacprowski, and Camilla Scheele. Multipep: a hierarchical deep learning approach for multi-label classification of peptide bioactivities. *Biology Methods and Protocols*, 6(1):bpab021, 2021.

Jiahui Guan, Peilin Xie, Dian Meng, Lantian Yao, Dan Yu, Ying-Chih Chiang, Tzong-Yi Lee, and Junwen Wang. Toxipep: Peptide toxicity prediction via fusion of context-aware representation and atomic-level graph. *Computational and Structural Biotechnology Journal*, 27:2347–2358, 2025.

Sudheer Gupta, Pallavi Kapoor, Kumardeep Chaudhary, Ankur Gautam, Rahul Kumar, and Gajendra PS Raghava. In silico approach for predicting toxicity of peptides and proteins. *PloS one*, 8(9):e73957–e73957, 2013.

Jiyun Han, Tongxin Kong, and Juntao Liu. Pepnet: an interpretable neural network for anti-inflammatory and antimicrobial peptides prediction using a pre-trained protein language model. *Communications Biology*, 7(1):1198, 2024.

Md Mehedi Hasan, Nalini Schaduangrat, Shaherin Basith, Gwang Lee, Watshara Shoombuatong, and Balachandran Manavalan. Hlppred-fuse: improved and robust prediction of hemolytic peptide and its activity by fusing multiple feature representation. *Bioinformatics*, 36(11):3350–3356, 2020.

Jesús Herrera-Bravo, Jorge G Farías, Fernanda Parraguez Contreras, Lisandra Herrera-Belén, and Jorge F Beltrán. Pep-predna+: A web server for prediction of highly specific peptides targeting voltage-gated na+ channels using machine learning techniques. *Computers in Biology and Medicine*, 145:105414, 2022.

Cataia Ives, Ivana Campia, Rong-Lin Wang, Clemens Wittwehr, and Stephen Edwards. Creating a structured adverse outcome pathway knowledgebase via ontology-based annotations. *Applied in vitro toxicology*, 3(4):298–311, 2017.

Aashish Jain and Daisuke Kihara. Nntox: gene ontology-based protein toxicity prediction using neural network. *Scientific reports*, 9 (1):17923, 2019.

Shihu Jiao, Xiucai Ye, Tetsuya Sakurai, Quan Zou, and Ruijun Liu. Integrated convolution and self-attention for improving peptide toxicity prediction. *Bioinformatics*, 40(5):btae297, 2024.

Shihu Jiao, Xiucai Ye, Tetsuya Sakurai, Quan Zou, Wu Han, and Chao Zhan. Integration of pre-trained protein language models with equivariant graph neural networks for peptide toxicity prediction. *BMC biology*, 23(1):229, 2025.

Xiaoyang Jing, Qiwen Dong, Daocheng Hong, and Ruqian Lu. Amino acid encoding methods for protein sequences: a comprehensive review and assessment. *IEEE/ACM transactions on computational biology and bioinformatics*, 17(6):1918–1931, 2019.

Dmitry A Karasev, Georgii S Malakhov, and Boris N Sobolev. Quantitative prediction of hemolytic activity of peptides. *Computational Toxicology*, 32:100335, 2024.

Hossein Khabbaz, Mohammad Hossein Karimi-Jafari, Ali Akbar Saboury, and Bagher BabaAli. Prediction of antimicrobial peptides toxicity based on their physico-chemical properties using machine learning techniques. *BMC bioinformatics*, 22(1):549, 2021.

KrenAI-Lab. BioSieve: Redundancy reduction and leakage-aware dataset partitioning for biological machine learning. `https://pypi.org/project/biosieve/`, 2026a. Python package, version 0.1.0, accessed 12 May 2026.

KrenAI-Lab. roxy. https://github.com/kren-ai-lab/roxy, 2026b. GitHub repository.

KrenAI-Lab. Sylphy: Protein sequence representation: encoders, embeddings, and reductions. `https://pypi.org/project/sylphy/`, 2026c. Python package, version 0.2.0, accessed 12 May 2026.

Ashok Kumar, Sonia Chadha, Mradul Sharma, and Mukesh Kumar. Deciphering optimal molecular determinants of non-hemolytic, cell-penetrating antimicrobial peptides through bioinformatics and random forest. *Briefings in Bioinformatics*, 26(1):bbaf049, 2025.

Yanru Li, Liming Gao, Wenqian Zhao, Ziyu Wang, Xianghan Xu, Jinwen Ji, Yuan Zhang, Yufeng Liu, Liangyun Zhang, Yong Yang, et al. Aipampds: an ai platform for antimicrobial peptide design and screening. *bioRxiv*, pages 2025–03, 2025.

Zeming Lin, Halil Akin, Roshan Rao, Brian Hie, Zhongkai Zhu, Wenting Lu, Nikita Smetanin, Allan dos Santos Costa, Maryam Fazel-Zarandi, Tom Sercu, Sal Candido, et al. Language models of protein sequences at the scale of evolution enable accurate structure prediction. *bioRxiv*, 2022.

Hongwu Lv, Ke Yan, and Bin Liu. Tppred-le: therapeutic peptide function prediction based on label embedding. *BMC biology*, 21(1): 238, 2023.

Tianyue Ma, Yanchao Liu, Bingxin Yu, Xin Sun, Huiyuan Yao, Chen Hao, Jianhui Li, Maryam Nawaz, Xun Jiang, Xingzhen Lao, et al. Dramp 4.0: an open-access data repository dedicated to the clinical translation of antimicrobial peptides. *Nucleic Acids Research*, 53(D1):D403–D410, 2025.

Charlene McQueen. *Comprehensive toxicology*. Elsevier, 2017.

Piotr Minkiewicz, Anna Iwaniak, and Małgorzata Darewicz. Biopep-uwm database of bioactive peptides: Current opportunities. *International journal of molecular sciences*, 20(23):5978, 2019.

Rajat Kumar Mondal, Debarup Sen, Ankish Arya, and Sintu Kumar Samanta. Developing anti-microbial peptide database version 1 to provide comprehensive and exhaustive resource of manually curated amps. *Scientific Reports*, 13(1):17843, 2023.

Vladimir Morozov, Carlos HM Rodrigues, and David B Ascher. Csm-toxin: a web-server for predicting protein toxicity. *Pharmaceutics*, 15(2):431, 2023.

Holly M Mortensen, Marvin Martens, Jonathan Senn, Trevor Levey, Chris T Evelo, Egon L Willighagen, and Thomas Exner. The aop-db rdf: applying fair principles to the semantic integration of aop data using the research description framework. *Frontiers in Toxicology*, 4:803983, 2022.

Raphaël Mourad. Mistral-prot-v1-134m (mistral for protein). `https://huggingface.co/RaphaelMourad/Mistral-Prot-v1-134M`, 2024. Hugging Face Model Repository.

Open Data Commons. Open data commons database contents license (DbCL) v1.0, 2008. URL `https://opendatacommons.org/licenses/dbcl/1-0/`. Consultado: 27 de julio de 2026.

Open Data Commons. Open data commons open database license (ODbL) v1.0, 2009. URL `https://opendatacommons.org/licenses/odbl/1-0/`. Consultado: 27 de julio de 2026.

Open Source Initiative. The MIT license, 2026. URL `https://opensource.org/licenses/MIT`. Consultado: 27 de julio de 2026.

AnQi Ouyang, Fang Ge, Jiangning Song, and Dong-Jun Yu. De novo design of functionally diverse and druggable antimicrobial peptides by diffusion and multimodal deep learning. *bioRxiv*, pages 2025–07, 2025.

Xiaoyong Pan, Jasper Zuallaert, Xi Wang, Hong-Bin Shen, Elda Posada Campos, Denys O Marushchak, and Wesley De Neve. Toxdl: deep learning using primary structure and domain embeddings for assessing protein toxicity. *Bioinformatics*, 36(21):5159–5168, 2020.

Gulnaz Perveen, Fahad Alturise, Tamim Alkhalifah, and Yaser Daanial Khan. Hemolytic-pred: a machine learning-based predictor for hemolytic proteins using position and composition-based features. *Digital Health*, 9:20552076231180739, 2023.

Fabien Plisson, Obed Ramírez-Sánchez, and Cristina Martínez-Hernández. Machine learning-guided discovery and design of non-hemolytic peptides. *Scientific reports*, 10(1):16581, 2020.

Peng Qiu, Hanqi Feng, Meng-Chun Zhang, and Barnabas Poczos. Amplyze: A deep learning model for predicting the hemolytic concentration. *arXiv preprint arXiv:2507.08162*, 2025.

Anand Singh Rathore, Akanksha Arora, Shubham Choudhury, Purva Tijare, and Gajendra PS Raghava. Toxinpred 3.0: An improved method for predicting the toxicity of peptides. *BioRxiv*, pages 2023–08, 2023.

Anand Singh Rathore, Saloni Jain, Shubham Choudhury, and Gajendra PS Raghava. Ntxpred2: a large language model for predicting neurotoxic peptides and neurotoxins. *bioRxiv*, 2025a.

Anand Singh Rathore, Nishant Kumar, Shubham Choudhury, Naman Kumar Mehta, and Gajendra PS Raghava. Prediction of hemolytic peptides and their hemolytic concentration. *Communications Biology*, 8(1):176, 2025b.

Sudipto Saha and Gajendra PS Raghava. Prediction of neurotoxins based on their function and source. *In silico biology*, 7(4-5):369–387, 2007.

Milad Salem, Arash Keshavarzi Arshadi, and Jiann Shiun Yuan. Ampdeep: hemolytic activity prediction of antimicrobial peptides using transfer learning. *BMC bioinformatics*, 23:1, 2022.

Harshika Sharma, Mayank Singh Thakur, Avinash Barala, Mohd Saad Khan, Sumit Bhagat, and Ganesh Bagler. Multitox: A sequence-based stacked ensemble model for multiclass protein toxin classification. *International Journal of Biological Macromolecules*, 327: 147399, 2025.

Neelam Sharma, Leimarembi Devi Naorem, Shipra Jain, and Gajendra PS Raghava. Toxinpred2: an improved method for predicting toxicity of proteins. *Briefings in bioinformatics*, 23(5):bbac174, 2022.

Hua Shi, Yan Li, Yi Chen, Yuming Qin, Yifan Tang, Xun Zhou, Ying Zhang, and Yun Wu. Toxmva: An end-to-end multi-view deep autoencoder method for protein toxicity prediction. *Computers in Biology and Medicine*, 151:106322, 2022.

Ayushi Singh, Kavin Raj SA, Anand Singh Rathore, and Gajendra P. S. Raghava. Hemolytik2: An updated database of hemolytic peptides and proteins. *bioRxiv*, 2025. doi: 10.1101/2025.05.12.653624. URL `https://www.biorxiv.org/content/early/2025/05/14/2025.05.12.653624`.

Sandeep Singh, Kumardeep Chaudhary, Sandeep Kumar Dhanda, Sherry Bhalla, Salman Sadullah Usmani, Ankur Gautam, Abhishek Tuknait, Piyush Agrawal, Deepika Mathur, and Gajendra PS Raghava. Satpdb: a database of structurally annotated therapeutic peptides. *Nucleic acids research*, 44(D1):D1119–D1126, 2016.

B Logan Spaller, Julie M Trieu, and Paulo F Almeida. Hemolytic activity of membrane-active peptides correlates with the thermodynamics of binding to 1-palmitoyl-2-oleoyl-sn-glycero-3-phosphocholine bilayers. *The Journal of membrane biology*, 246(3):257–262, 2013.

Sebastian Spänig, Siba Mohsen, Georges Hattab, Anne-Christin Hauschild, and Dominik Heider. A large-scale comparative study on peptide encodings for biomedical classification. *NAR genomics and bioinformatics*, 3(2):lqab039, 2021.

Zhelyazko Terziyski, Margarita Terziyska, Ivelina Deseva, Stanka Hadzhikoleva, Albert Krastanov, Dasha Mihaylova, and Emil Hadzhikolev. Peplab platform: Database and software tools for analysis of food-derived bioactive peptides. *Applied Sciences*, 13(2):961, 2023.

Patrick Brendan Timmons and Chandralal M Hewage. Happenn is a novel tool for hemolytic activity prediction for therapeutic peptides which employs neural networks. *Scientific reports*, 10(1):10869, 2020.

Andrey A Toropov, Alla P Toropova, Valentin O Kudyshkin, Emilio Benfenati, Danuta Leszczynska, and Jerzy Leszczynski. Simulation peptide toxicity using the fragments of local symmetry in amino acid sequences. *BioSystems*, page 105554, 2025.

Duong Thanh Tran, Nhat Truong Pham, Nguyen Doan Hieu Nguyen, Leyi Wei, and Balachandran Manavalan. Hypeptox-fuse: An interpretable hybrid framework for accurate peptide toxicity prediction fusing protein language model-based embeddings with conventional descriptors. *Journal of Pharmaceutical Analysis*, page 101410, 2025.

Hao Wan, Qing Liu, and Ying Ju. Utilize a few features to classify presynaptic and postsynaptic neurotoxins. *Computers in Biology and Medicine*, 152:106380, 2023.

Feng Wang, Peilin Xie, Xingqiao Lin, Jiahui Guan, Chang Liu, Xuxin He, Tzong-Yi Lee, Leyi Wei, Xiangrong Liu, and Lantian Yao. Bitoxnet: a deep learning framework integrating multimodal features for accurate identification of neurotoxic peptides and proteins. *BMC biology*, 24(1):89, 2026.

Jen-Hung Wang and Ting-Yi Sung. Toxteller: predicting peptide toxicity using four different machine learning approaches. *ACS omega*, 9(29):32116–32123, 2024.

Li Wang, Xiangzheng Fu, Jiahao Yang, Xinyi Zhang, Xiucai Ye, Yiping Liu, Tetsuya Sakurai, and Xiangxiang Zeng. Moformer: Multi-objective antimicrobial peptide generation based on conditional transformer joint multi-modal fusion descriptor. *arXiv preprint arXiv:2406.02610*, 2024a.

Li Wang, Yiping Liu, Xiangzheng Fu, Xiucai Ye, Junfeng Shi, Gary G Yen, Quan Zou, Xiangxiang Zeng, and Dongsheng Cao. Hmamp: designing highly potent antimicrobial peptides using a hypervolume-driven multiobjective deep generative model. *Journal of Medicinal Chemistry*, 68(8):8346–8360, 2025a.

Yanling Wang, Na Li, Xiao Wang, Feng Cao, Shuwen Xiong, and Leyi Wei. Integrating protein language models and geometric deep learning for peptide toxicity prediction. *Journal of Chemical Information and Modeling*, 65(14):7800–7810, 2025b.

Yue Wang, Haifan Gong, Xiaojuan Li, Lixiang Li, Yinuo Zhao, Peijing Bao, Qingzhou Kong, Boyao Wan, Yumeng Zhang, Jinghui Zhang, et al. De novo multi-mechanism antimicrobial peptide design via multimodal deep learning. *BioRxiv*, pages 2024–01, 2024b.

Lesong Wei, Xiucai Ye, Yuyang Xue, Tetsuya Sakurai, and Leyi Wei. Atse: a peptide toxicity predictor by exploiting structural and evolutionary information based on graph neural network and attention mechanism. *Briefings in bioinformatics*, 22(5):bbab041, 2021.

Lesong Wei, Xiucai Ye, Tetsuya Sakurai, Zengchao Mu, and Leyi Wei. Toxibtl: prediction of peptide toxicity based on information bottleneck and transfer learning. *Bioinformatics*, 38(6):1514–1524, 2022.

William C Wimley. Describing the mechanism of antimicrobial peptide action with the interfacial activity model. *ACS chemical biology*, 5(10):905–917, 2010.

Thet Su Win, Aijaz Ahmad Malik, Virapong Prachayasittikul, Jarl E S Wikberg, Chanin Nantasenamat, and Watshara Shoombuatong. Hemopred: a web server for predicting the hemolytic activity of peptides. *Future medicinal chemistry*, 9(3):275–291, 2017.

Peilin Xie, Lantian Yao, Jiahui Guan, Chia-Ru Chung, Zhihao Zhao, Feiyu Long, Zhenglong Sun, Tzong-Yi Lee, and Ying-Chih Chiang. Consamphemo: A computational framework for predicting hemolysis of antimicrobial peptides based on machine learning approaches. *Protein Science*, 34(7):e70087, 2025.

Jing Xu, Fuyi Li, Chen Li, Xudong Guo, Cornelia Landersdorfer, Hsin-Hui Shen, Anton Y Peleg, Jian Li, Seiya Imoto, Jianhua Yao, et al. iampcn: a deep-learning approach for identifying antimicrobial peptides and their functional activities. *Briefings in Bioinformatics*, 24(4):bbad240, 2023.

Sen Yang and Piao Xu. Hemodl: Hemolytic peptides prediction by double ensemble engines from rich sequence-derived and transformer-enhanced information. *Analytical Biochemistry*, 690:115523, 2024.

Yang Yang, Haohan Zhang, and Mauno Vihinen. Protoxin, a predictor of protein toxicity. *Toxins*, 17(10):489, 2025.

Adiba Yaseen, Sadaf Gull, Naeem Akhtar, Imran Amin, and Fayyaz Minhas. Hemonet: Predicting hemolytic activity of peptides with integrated feature learning. *Journal of Bioinformatics and Computational Biology*, 19(05):2150021, 2021. doi: 10.1142/S0219720021500219. URL `https://doi.org/10.1142/S0219720021500219`. PMID: 34353244.

Qiule Yu, Zhixing Zhang, Guixia Liu, Weihua Li, and Yun Tang. Toxgin: an in silico prediction model for peptide toxicity via graph isomorphism networks integrating peptide sequence and structure information. *Briefings in Bioinformatics*, 25(6):bbae583, 2024.

Shengli Zhang, Jingyi Ren, and Yunyun Liang. An innovative peptide toxicity prediction model based on multi-scale convolutional neural network and residual connection. *Bioinformatics*, 41(10):btaf462, 2025.

Ya Zhao, Shengli Zhang, and Yunyun Liang. Hemofuse: multi-feature fusion based on multi-head cross-attention for identification of hemolytic peptides. *Scientific Reports*, 14(1):22518, 2024.

Yuhong Zhao, Shijing Wang, Wenyi Fei, Yuqi Feng, Le Shen, Xinyu Yang, Min Wang, and Min Wu. Prediction of anticancer peptides with high efficacy and low toxicity by hybrid model based on 3d structure of peptides. *International journal of molecular sciences*, 22(11):5630, 2021.

Lin Zhu, Yi Fang, Shuting Liu, Hong-Bin Shen, Wesley De Neve, and Xiaoyong Pan. Toxdl 2.0: Protein toxicity prediction using a pretrained language model and graph neural networks. *Computational and Structural Biotechnology Journal*, 27:1538–1549, 2025.

Wen Zhu, Yuxin Guo, and Quan Zou. Prediction of presynaptic and postsynaptic neurotoxins based on feature extraction. *Math Biosci Eng*, 18(5):5943–58, 2021.

Shengxin Zhuang, John Tanner, Yusen Wu, Du Huynh, Wei Liu, Xavier Cadet, Nicolas Fontaine, Philippe Charton, Cedric Damour, Frederic Cadet, et al. Non-hemolytic peptide classification using a quantum support vector machine: S. zhuang et al. *Quantum Information Processing*, 23(11):379, 2024.
